# The emergence of successful *Streptococcus pyogenes* lineages through convergent pathways of capsule loss and recombination directing high toxin expression

**DOI:** 10.1101/684167

**Authors:** Claire E. Turner, Matthew T. G. Holden, Beth Blane, Carolyne Horner, Sharon J. Peacock, Shiranee Sriskandan

## Abstract

Gene transfer and homologous recombination in *Streptococcus pyogenes* has the potential to trigger the emergence of pandemic lineages, as exemplified by lineages of *emm*1 and *emm*89 that emerged in the 1980s and 2000s respectively. Although near-identical replacement gene transfer events in the *nga* (NADase) and *slo* (Streptolysin O) locus conferring high expression of these toxins underpinned the success of these lineages, extension to other *emm*-genotype lineages is unreported. The emergent *emm*89 lineage was characterised by five regions of homologous recombination additional to *nga/slo*, including complete loss of the hyaluronic acid capsule synthesis locus *hasABC,* a genetic trait replicated in two other leading *emm* types and recapitulated by other *emm* types by inactivating mutations. We hypothesised that other leading genotypes may have undergone a similar recombination events. We analysed a longitudinal dataset of genomes from 344 clinical invasive disease isolates representative of locations across England, dating from 2001 to 2011, and an international collection of *S. pyogenes* genomes representing 54 different genotypes, and found frequent evidence of recombination events at the *nga*-*slo* locus predicted to confer higher toxin expression. We identified multiple associations between recombination at this locus and inactivating mutations within *hasA/B,* suggesting convergent evolutionary pathways in successful genotypes. This included common genotypes *emm*28 and *emm*87. The combination of no or low capsule, and high expression of *nga* and *slo,* may underpin the success for many emergent *S. pyogenes* lineages of different genotypes, triggering new pandemics and could change the way *S. pyogenes* causes disease.

**Importance:** *Streptococcus pyogenes* is a genetically diverse pathogen, with over 200 different genotypes defined by *emm* typing, but only a minority of these genotypes are responsible for majority of human infection in high income countries. Two prevalent genotypes associated with disease rose to international dominance following recombination of a toxin locus that conferred increased expression. Here, we found that recombination of this locus and promoter has occurred in other diverse genotypes, events that may allow these genotypes to expand in the population. We identified an association between the loss of hyaluronic acid capsule synthesis and high toxin expression, which we propose may be associated with an adaptive advantage. As *S. pyogenes* pathogenesis depends both on capsule and toxin production, new variants with altered expression may result in abrupt changes in the molecular epidemiology of this pathogen in the human population over time.

## Introduction

The capacity for the bacterial human pathogen *Streptococcus pyogenes* to undergo genetic exchange, independent of known bacteriophages or mobile elements, is not well understood, yet recent evidence suggests it underpins the emergence of successful new variants that rapidly rise to international dominance. Homologous recombination of a chromosomal region encompassing the toxin genes *nga* (encoding for NADase), *ifs* (encoding the inhibitor for NADase) and *slo* (encoding for Streptolysin O), which was dated to have occurred in the mid-1980s, is thought to have driven the rise of *emm*1 to almost global dominance (1). The homologous recombination event resulted in increased *nga*/*slo* expression compared to the previous variant, linked to the gain of a highly active *nga/ifs/slo* promoter in the new *emm*1 variant compared to the previous variant (2).

A very similar recombination event was recently identified in the genotype *emm*89. A new variant of *emm*89 sequence type (ST) 101 (also referred to as Clade 3) emerged, having undergone six regions of predicted homologous recombination compared to its ST101 predecessor (also referred to as Clade 2) (3, 4). One of the six regions encompassed the *nga/ifs/slo* locus, comprising a region almost identical to *emm*1, that conferred similarly high expression of *nga* and *slo* compared to the previous variant. Another recombination region within the emergent ST101 *emm*89 resulted in the loss of the hyaluronic acid capsule. We dated the emergence of this new acapsular, high toxin expressing ST101 *emm*89 lineage to the mid-1990s, but there was a rapid increase and rise to dominance in the UK between 2005-2010 (3). The lineage is now the dominant form of *emm*89 in the UK as well as other parts of the world including Europe, North America and Japan (4–8).

Given that recombination associated with *nga*/*ifs*/*slo* can give rise to new successful *S. pyogenes* variants, we hypothesised that this may be a feature common to other successful *emm-*types. To determine if this is the case, we sequenced the genomes of 344 *S. pyogenes* invasive disease isolates originating from hospitals across England between 2001-2011, and compared the data with other available historical and contemporary international *S. pyogenes* whole genome sequence (WGS) data. We identified that recombination of the *nga-ifs-slo* locus has occurred in other leading *emm*-types, supporting the hypothesis that it can underpin the emergence and success of new lineages. We also identified an association of *nga-ifs-slo* recombination towards a high activity promoter variant with inactivating mutations within the capsule locus. This suggests that loss of capsule may also provide an advantage to certain genotypes, either through a direct effect on pathogenesis or an association with the process of recombination.

## Results

### Genetic characterisation of bacteraemia isolates

We performed whole genome sequencing of 344 *S. pyogenes* invasive isolates collected from hospitals across England by the British Society for Antimicrobial Chemotherapy (BSAC) Bacteraemia Resistance Surveillance Programme during 2001-2011. Forty-four different *emm*-types were identified from *de novo* assembly, with the most common being *emm*1 (n=64, 18.6%), *emm*12 (n=34, 9.9%), *emm*89 (n=32, 9.3%), *emm*3 (n=28, 8.1%), *emm*87 (n=22, 6.4%) and *emm*28 (n=15, 4.4%) (Supplementary Figure 1). Antimicrobial susceptibilities were typical for *S. pyogenes* with 100% isolates susceptible to penicillin, and 20% resistant to macrolides; detailed susceptibilities and associated genotypes are reported in Supplementary Table 1.

The phylogenetic distribution of the 344 isolates based on core genome variation revealed distinct clustering by *emm*-type, each forming single lineages with the exceptions of *emm*44, *emm*90 and *emm*101, each of which formed two lineages (Figure 1A). Pairwise distances between isolates gave a median of just 45 SNPs separating the genomes of isolates of the same *emm*-genotype (range 0-15,137 SNPs), compared to a median of 15,648 SNPs separating the genomes of isolates of different *emm*-types (range 5312-18,317 SNPs) (Figure 1B). The genotypes *emm*44, *emm*90 and *emm*101 gave the highest SNP distance for the intra-*emm* comparison (13,494 - 15,137 SNPs) which approaches the median level observed between *emm*-types. This indicated that while other genotypes represent a relatively conserved chromosomal genetic background, the populations of *emm*44, *emm*90 and *emm*101 exhibit more diverse chromosomal backgrounds despite representing the same *emm*-type, potentially due to *emm* gene switching.

**Figure 1.**
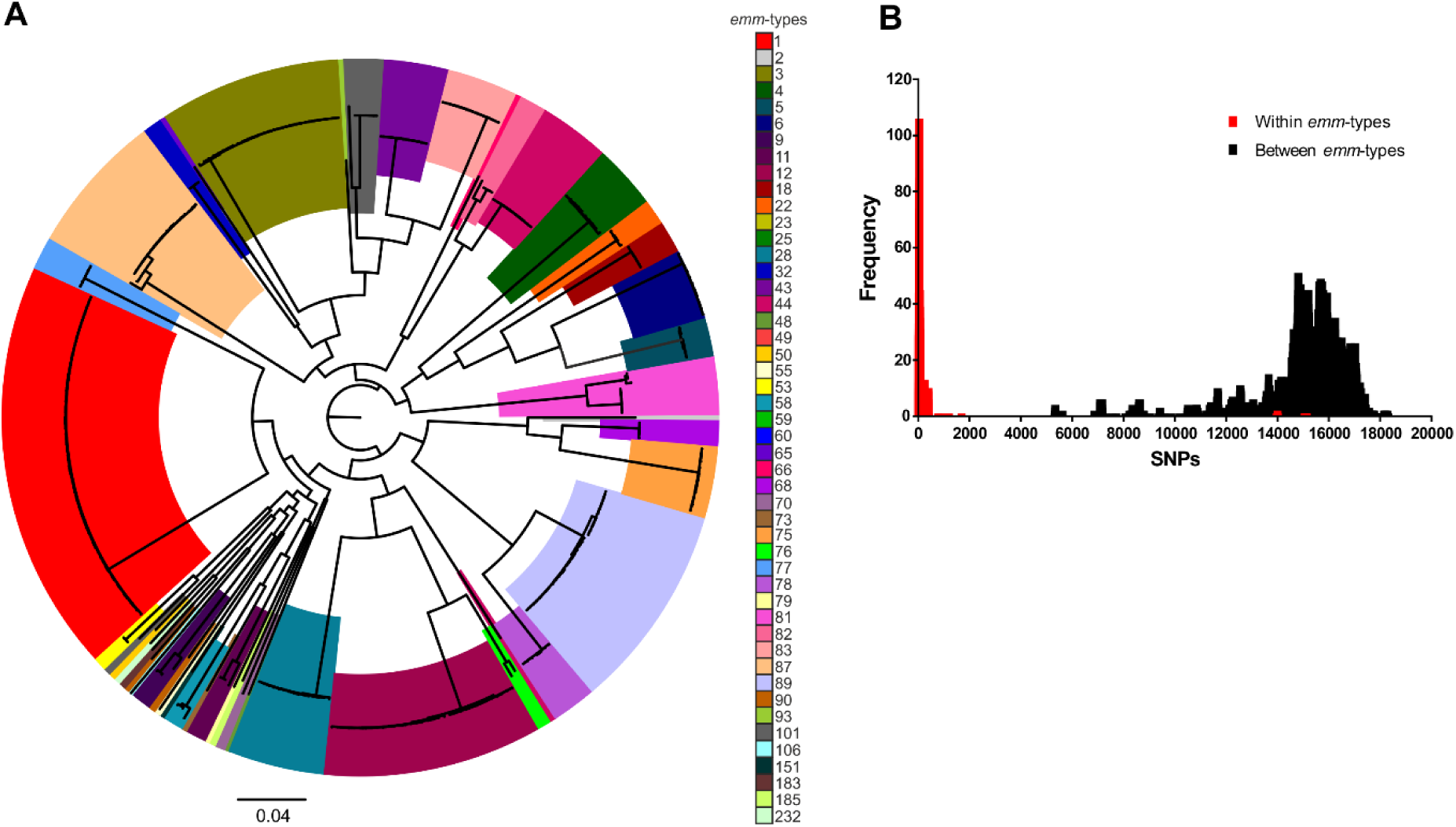
Low diversity within *emm* genotypes. (**A**) A maximum likelihood phylogenetic tree constructed from core SNPs extracted after mapping all 344 BSAC isolates to the complete reference strain H293, identified that the majority of isolates cluster by *emm* genotype. Exceptions were *emm*44, *emm*90 and *emm*101, each of which were present as two separate lineages. (**B**) As reflected by the phylogenetic tree, the number of SNPs separating isolates was high (>5000) when the genomes of isolates of different *emm*-types were compared (black bars). This was much lower when comparisons were made between the genomes of isolates of the same *emm*-type (red bars).

### High level of variation within the nga-ifs-slo locus

In order to identify the level of variation within the *nga-ifs-slo* locus we extracted the sequence from the 3’ end of *nusG* (immediately upstream of *nga*) to the 3’ end of *slo* (P-*nga-ifs-slo*), comprising the entire locus and all upstream sequence including the predicted ∼67bp *nga/ifs/slo* promoter region (9). We constructed a phylogenetic tree from SNPs within P-*nga-ifs-slo* region and compared it to the phylogeny constructed with SNPs extracted from a whole genome comparison to a reference *emm*89 genome, H293 (Figure 2). In most cases, a single unique P-*nga-ifs-slo* variant was associated with each *emm* genotype, consistent with a conserved chromosome. The main exception to this was the P-*nga-ifs-slo* variant found in modern (post 1980s M1T1) *emm*1, which was also found in all *emm*12, all *emm*22 (a lineage known to be acapsular), and 11 of the 32 *emm*89 isolates. These 11 *emm*89 represented the emergent acapsular ST101 variant, whilst the remaining 21 *emm*89 isolates represented the original encapsulated ST101 variant, with a different unique P-*nga-ifs-slo* as previously reported (3). The entire *emm*75 population and one of the two *emm*76 isolates were associated with a P-*nga-ifs-slo* variant that was closely related to the *emm*1-like variant. All but two *emm*87 isolates had a P-*nga-ifs-slo* variant also found in the acapsular lineage *emm*4 (Figure 2). The presence of multiple P-*nga-ifs-slo* variants within single *emm* genotypes, where the core chromosome was otherwise relatively conserved, indicated that gene transfer and recombination are responsible for the variation rather than extensive genome-wide divergence or *emm* ‘switching’.

**Figure 2.**
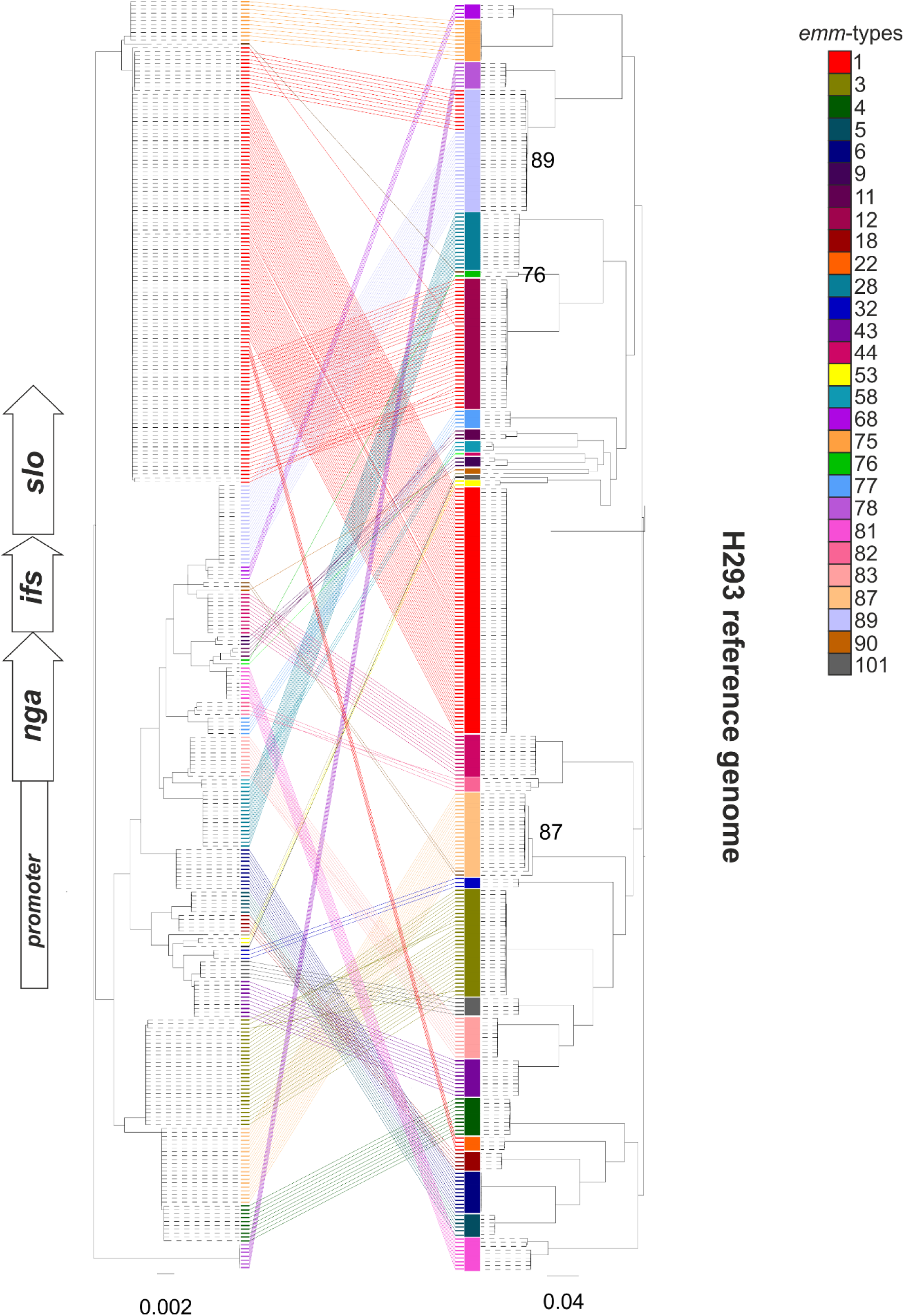
Comparison of the variation within the P-*nga-ifs-slo* region and core chromosome. A maximum likelihood phylogenetic tree was constructed from SNPs extracted from an alignment of the *nga-ifs-slo* locus and associated upstream region to include the promoter (P-*nga-ifs-slo*) extracted from *de novo* assemblies of 344 BSAC *S. pyogenes* (Left tree). This was compared to the phylogenetic tree constructed using SNPs across the entire genome after mapping to the H293 reference genome (Right tree). Only *emm* genotypes represented by two or more isolates were included. Coloured blocks on the right tree represent *emm*-type. Variants of the P-*nga-ifs*-*slo* are of the same colour if unique to that genotype. The P-*nga-ifs-slo* variant found in *emm*1 (red) was common to other genotypes *emm*12, *emm*22 and some *emm*89. The genotypes *emm*76, *emm*87 and *emm*89 are indicated as they are linked to more than one variant of P-*nga-ifs-slo*. Scale bar represents substitution per site.

### Variants of the nga-ifs-slo promoter associated with altered expression

Recombination of P-*nga-ifs-slo* and surrounding regions in *emm*1 and *emm*89 conferred higher activity and expression of NGA and SLO (1, 3, 10). This change in expression was linked to the combination of three key residues at −27, −22 and −18 within the *nga-ifs-slo* promoter. A_−27_G_−22_T_−18_ at these key sites was associated with high *nga-ifs-slo* promoter activity in *emm*1 and emergent *emm*89 following recombination (also referred to as Pnga3) compared to low promoter activity of historical *emm*1 and *emm*89, associated with the key site combinations A_−27_T_−22_C_−18_ and G_−27_T_−22_T_−18_ respectively (2) (Figure 3A). We compared the ∼67bp *nga-ifs-slo* promoter region of the 344 BSAC collection isolate genomes to identify different variants. We expanded the data analysed by including assembled genome data from over 5000 isolates representing 54 different *emm* types: from Cambridge University Hospital (CUH) (11), the rest of England and Wales collected by Public Health England (PHE) in 2014/2015 (PHE-2014/15) (12, 13) and from the USA collected by the Active Bacterial Core Surveillance System (ABCs) in 2015 (ABCs-2015) (14). We excluded 39 *emm*-types represented by fewer than 3 isolates (Supplementary Table 2).

**Figure 3.**
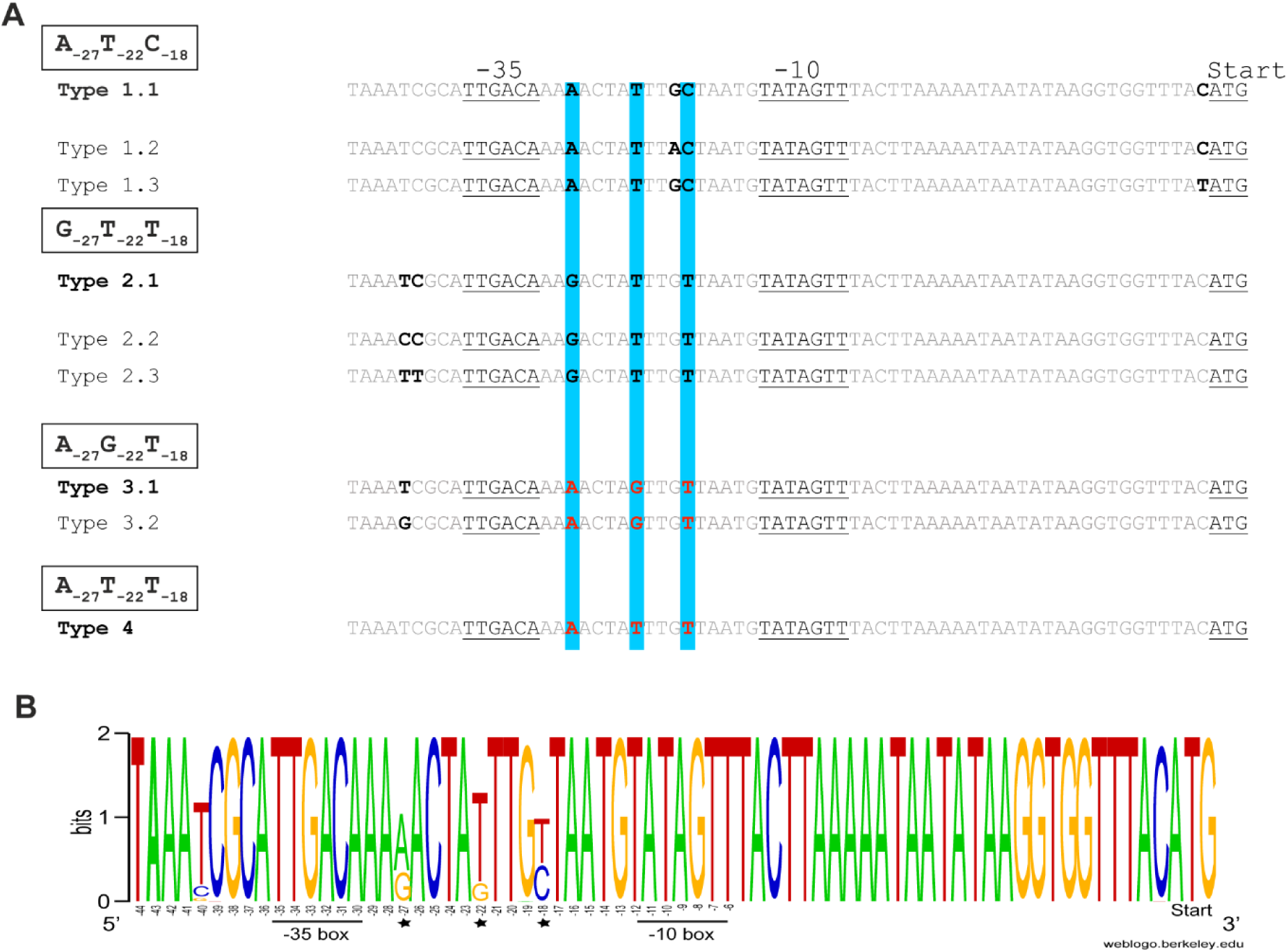
Variants of the *nga-ifs-slo* promoter. **(A)** The three key residues predicted to influence promoter activity are highlighted blue with those associated with high activity in red. We identified four combinations of these residues (four promoter types) with subtype variants differing in residues other than −27, −22 and −18 (residue positions relative to the underlined −35 and −10 regions) in the predicted 67bp promoter region (9). The combination of A_−27_T_−22_C_−18_ subtype 1.1 in historical *emm*1 and G_−27_T_−22_T_−18_ subtype 2.1 in older *emm*89 have been shown to be associated with low level promoter activity. A_−27_G_−22_T_−18_ subtype 3.1 promoter in modern *emm*1 and emergent variant *emm*89 has been shown to have high activity. A_−27_T_−22_T_−18_ subtype 4 promoter has also been shown to have high activity in *emm*28 (15). Subtypes 1.2, 1.3 and 2.3 were restricted to *emm*9, *emm*88 and *emm*32 strains respectively. **(B)** Weblogo representation of the variability in the 67bp promoter region of *nga/ifs/slo* within the 54 different *emm*-types. Key residues −27, −22, −18 are highlighted (star) and their positions are relative to the −35 and −10 boxes. Figure generated using weblogo.berkeley.edu.

Four combinations of the −27, −22 and −18 residues were found across all 5271 isolates (Table 1); variant 1 A_−27_T_−22_C_−18_ and variant 2 G_−27_T_−22_T_−18_ are associated with low promoter activity, while variant 3 A_−27_G_−22_T_−18_ and variant 4 A_−27_T_−22_T_−18_ are associated with high promoter activity. We also identified subtypes of the 67bp promoter region which varied at bases other than −27, −22 and −18 (Figure 3A and B, Table 1). A_−27_T_−22_C_−18_ variant subtype 1.1 and G_−27_T_−22_T_−18_ variant subtype 2.1 have both previously been confirmed to have low promoter activity (2) and were the most common variants found across genotypes. Other subtypes of these variants were restricted to single genotypes except G_−27_T_−22_T_−18_ variant subtype 2.2, which differed by a single substitution of C for a T residue at −40bp. Two subtypes of the high activity variant A_−27_G_−22_T_−18_ were found, the most common being subtype 3.1 associated with *emm*1 and emergent *emm*89, and subtype 3.2 which was found predominantly in the genomes of *emm*4 and *emm*87 and which differed from subtype 3.1 by a single substitution of G for T at −40bp. We measured the activity of NADase in the culture supernatant of strains representing different promoter subtypes and predict that the presence of T/G/C at −40bp does not affect activity of the promoter (Supplementary Figure 3). The fourth promoter variant, A_−27_T_−22_T_−18_ is also associated with high activity (15) and was identified in the genomes of *emm*28, *emm*75 and all *emm*78. Only three *emm*-types were exclusively associated with the high activity promoter variant A_−27_G_−22_T_−18_; *emm*1, *emm*3 and *emm*12. Other *emm*-types with the high activity promoter variant also had one or more of the other three promoter variants, suggesting a mixed population or, as in the case of *emm*89, an evolving population.

**Table 1.** Three key residue variants within the nga-ifs-slo promoter.

| Promoter variant | Type | Genotype (% of isolates) |
| --- | --- | --- |
| A-27T-22C-18 | 1.1 | **4 (1)\***, 8 (100), **9 (92)**, 11 (100), **22 (3)**, **25 (33)**, **28 (87.7)**, 33 (100), 41 (100), 43 (100), **44 (9)**, 49 (100), 53 (100), **58 (15)**, 60 (100), 63 (100), **75 (9)**, **76 (41)**, **77 (29)**, **81 (23)**, **82 (1)**, **88 (33)**, **89 (1)**, **90 (4)**, 92 (100), **94 (6)**, 101 (100), **102 (50)**, **103 (17)**, 106 (100), **108 (89)**, 110 (100), 113 (100), 151 (100), 168 (100), 232 (100) |
|  | 1.2 | 9(8) |
|  | 1.3 | 88(67) |
| G-27T-22T-18 | 2.1 | 5 (100), 6 (100), 18 (100), **25 (67)**, **44 (28)**, 68 (100), **75 (1)**, **76 (5)**, **77 (1)**, **82 (1)**, **87 (2)**, **89 (6)**, **90 (96)**, 91 (100), **102 (50)**, **103 (83)**, 104 (100), 118 (100) |
|  | 2.2 | 2 (100), 27 (100), **44 (62)**, **58 (85)**, 59 (100), 73 (100), **76 (11)**, **77 (36)**, **82 (89)**, 83 (100) |
|  | 2.3 | 32 (100) |
| A-27G-22T-18 | 3.1 | 1(100), 3 (100), 12 (100), **22 (97)**, **75 (90)**, **76 (43)**, **81 (77)**, **82 (9)**, **89 (93)**, **94 (94)**, **108 (11)** |
|  | 3.2 | **4 (99)**, **28 (0.3)**, **77 (34)**, **87 (98)** |
| A-27T-22T-18 | 4 | **28 (12)**, 78 (100) |

\* genotypes in bold have more than one variant within the population

To identify any possible recombination events surrounding the P-*nga-ifs-slo* region, we generated a maximum likelihood phylogeny based on SNPs within the P-*nga-ifs-slo* (Figure 4). This identified several more instances where more than one variant was associated with a single genotype and a cluster of variants with high activity associated promoter residues A_−27_G_−22_T_−18_.

**Figure 4.**
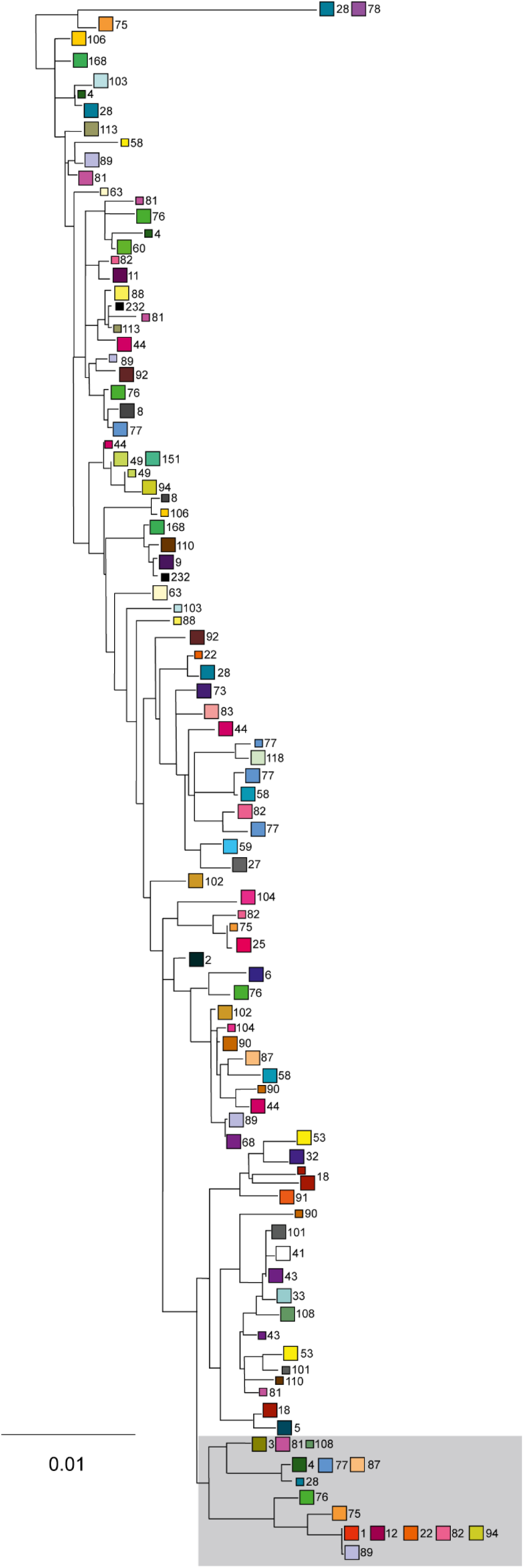
Variants of P-*nga-ifs-slo* within *emm* genotypes. The P-*nga-ifs-slo* was extracted from 5271 assembled genomes and aligned for SNP extraction, and these used for maximum likelihood midpoint-rooted phylogenetic tree construction. Squares represent multiple identical sequences (larger squares) or single sequences (smaller squares) of individual *emm*-types. The *emm*-type is given next to each square. Shaded region; high activity cluster. Scale represents substitutions per sites.

We sought evidence for acquisition of the high activity-associated promoter A_−27_G_−22_T_−18_ variant by *emm* genotypes where the dominant or ancestral state was a low activity-associated promoter; these included, in addition to the aforementioned *emm*89: *emm*75, *emm*76, *emm*77, *emm*81, *emm*82, *emm*87, *emm*94 and *emm*108, all of which are *emm* types frequently identified in the UK and the USA (12–14). Although one *emm*28 was found to carry the high activity-associated promoter, the rest of the *emm*28 population was divided between either A_−27_T_−22_C_−18_ or A_−27_T_−22_T_−18_ variants. The data pointed to a switch in P-*nga-ifs-slo* in all cases rather than an *emm* switch, except for *emm*82, where the *emm*82 gene has replaced the *emm*12 gene in an *emm*12 genetic background (14).

### High level of mutations within the capsule locus leading to truncations of HasA or HasB

As well as recombination around the P-*nga-ifs-slo* region, the emergent ST101 variant of *emm*89 had also undergone recombination surrounding the *hasABC* locus, and, in place of the *hasABC* genes, was a region of 156bp that was not found in genotypes with the capsule locus but is found in the acapsular *emm*4 and *emm*22 isolates (3). To identify any similar events in other genotypes, we examined the sequences of *hasA*, *hasB,* and *hasC* in the assemblies of isolates from the BSAC collection as well as CUH (11), PHE-2014/15 and ABCs-2015 collections for gene presence as well as premature stop codon mutations or missing genes (Figure 5). The *hasABC* locus was absent in the majority of *emm*89 isolates, consistent with the previous observations describing the recent emergence of the acapsular *emm*89 variant (3). Similarly, the *hasABC* genes were absent in all *emm*4 and *emm*22 isolates, as previously identified (16), except for two *emm*4 isolates and one *emm*22 isolate which had an intact *hasABC* locus predicted to encode full length proteins. We confirmed the genotypes of these isolates by *emm*-typing the assembled genomes; MLST and phylogenetic analysis indicated they both had a very different genetic background to other *emm*4 or *emm*22 populations suggesting these were not typical of these *emm* types, and therefore they represent examples of *emm* switching. Interestingly, we also identified a similar replacement of *hasABC* for the 156bp region in one *emm*28 isolate (PHE-2014/15, GASEMM1261 (13)), but phylogenetic analysis suggested this was highly divergent to the rest of the *emm*28 population, likely to represent another example of *emm* switching. Isolated examples of individual *hasA* or *hasB* gene loss were identified in the genomes of isolates belonging to *emm*1 (n=1), *emm*3 (n=1), *emm*11 (n=1), *emm*12 (n=4) and *emm*108 (n=2).

**Figure 5.**
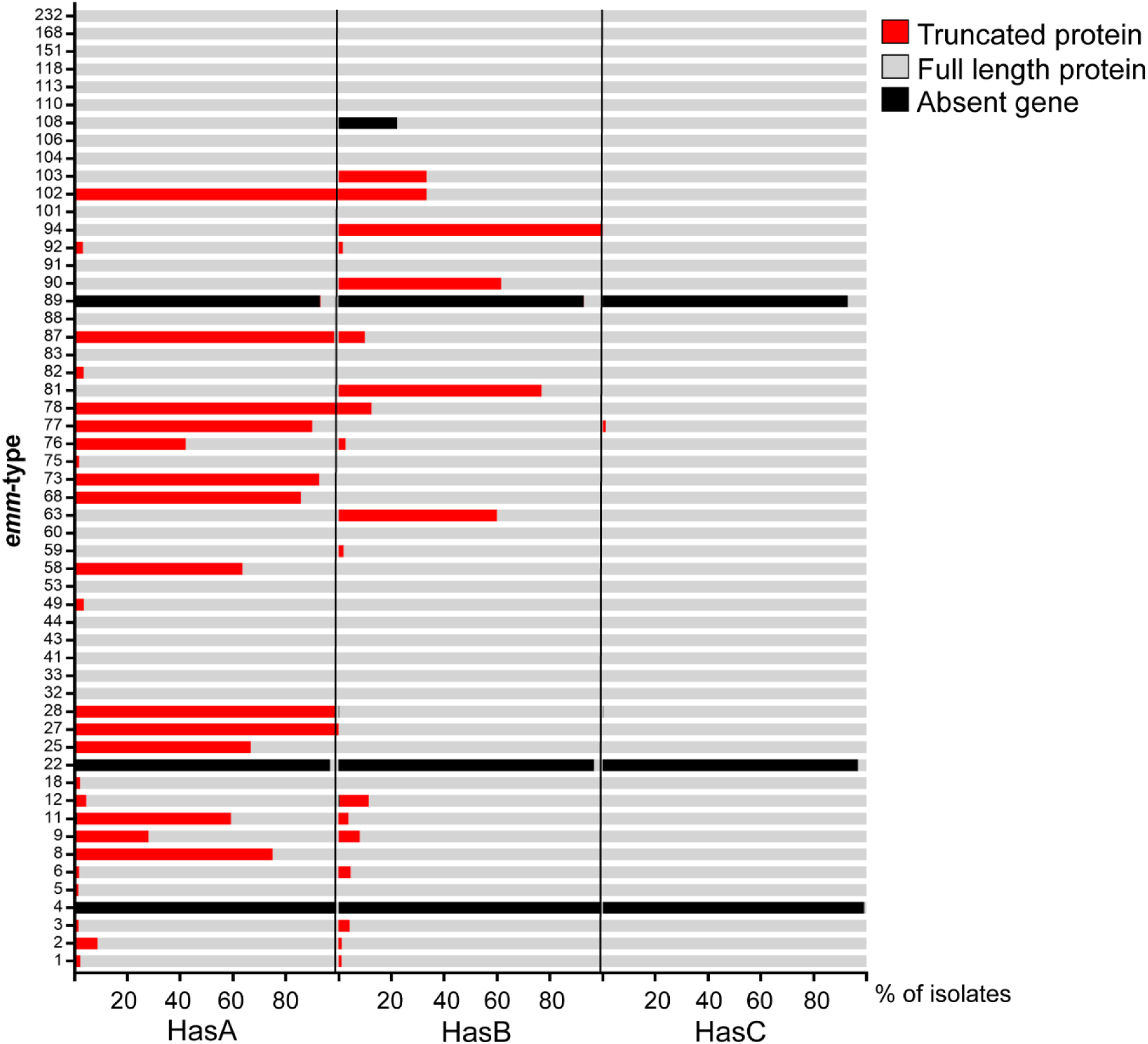
Non-functional mutations within the capsule locus genes. The *hasABC* genes were extracted from the assembled genomes of BSAC, CUH, PHE-2015/15, and ABCs-2015 isolate collections and polymorphisms or indels leading to nonsense mutations and premature stop codons were identified, as well as gene absence. The percentage of isolates with full length (grey), truncated (red) or absent (black) HasA, HasB or HasC is depicted for each of the 54 *emm*-types. *emm*-types with fewer than 3 isolates were excluded. N = 5271 isolates shown. Mutations in *hasA* were detected in more than 50% of isolates belonging to genotypes *emm*8 (n=3/4), *emm*11 (n=63/108), *emm*25 (n=2/3), *emm*27 (n=3/3), *emm*28 (n=358/363), *emm*58 (n=21/33), *emm*68 (n=12/14), *emm*73 (n=25/27), *emm*77 (n=72/80), *emm*78 (n=8/8), *emm*87 (n=119/121) and *emm*102 (n=6/6). Mutations in *hasB* were detected in 100% of *emm*94 isolates (n=54/54) and 60-77% of *emm*63 (n=3/5), *emm*81 (n=50/65) and *emm*90 (n=16/26) isolates.

The majority of genotypes (n=35/54, 65%) had isolates without genes or truncation mutations in at least one of *hasABC* genes. In some cases, a consistent mutation could be identified across the genotype (Figure 5). Mutations in *hasC* were rare and only detected in one isolate, an *emm*77 which also had a mutation within *hasA*. Within seven of the eight *emm*-types for which we identified potential P-*nga-ifs-slo* recombination, a high percentage of isolates had inactivating mutations *hasA* and *hasB* suggesting a possible association between an acapsular and recombination of P-*nga-ifs-slo*.

### Recombination of P-nga-ifs-slo and surrounding regions

To confirm our prediction that genotypes *emm*28, *emm*75, *emm*76, *emm*77, *emm*81, *emm*87, *emm*94 and *emm*108 had undergone recombination around P-*nga-ifs-slo*, we mapped all the genome sequence data for each genotype to the *emm*89 reference genome H293. Gubbins analysis of SNP clustering predicted regions of recombination spanning the *nga*-*ifs-slo* region and varying in length in all eight genotypes (Figure 6). To analyse recombination of these genotypes and potential capsule loss further, we studied the population structure of each genotype individually.

**Figure 6.**
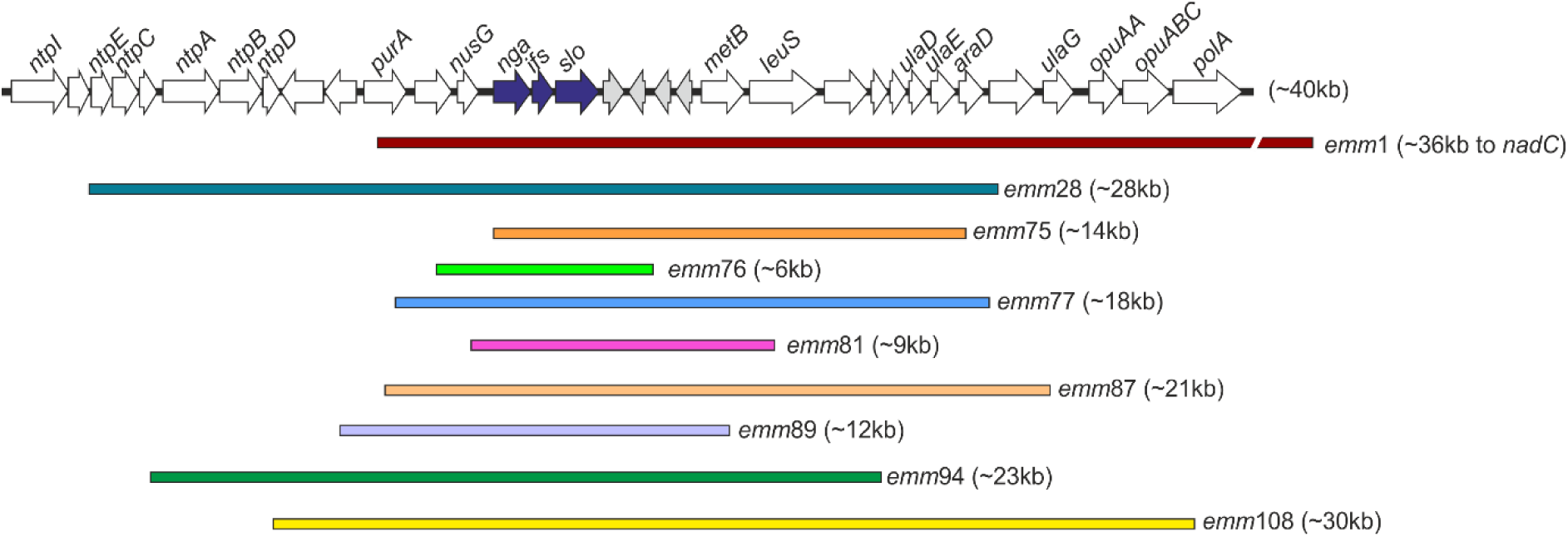
Regions of recombination spanning the P-*nga-ifs-slo* locus. Recombination across the *nga, ifs* and *slo* genes (blue arrows) was identified in eight genotypes in addition to the previously described *emm*1 and *emm*89. Length of recombination, predicted by SNP cluster analysis, ranged from ∼6kb to 36kb. With the exception of *emm*75, all regions also encompassed the promoter of *nga-ifs-slo*. All regions are shown relative to a ∼40kb region within the reference genome H293 and genes within this region are depicted as arrows. Recombination in *emm*1 extended beyond that depicted here and is shown as a broken line.

### Recombination within emm28 and emm87 around P-nga-ifs-slo and the capsule locus

The genotypes *emm*28 and *emm*87 were the sixth and fifth most common in the BSAC collection, and *emm*28 has previously be noted to be a major cause of infection in high income countries (17). We focussed attention on *emm*28 and *emm*87 as there has been little genomic work on these genotypes so far.

All BSAC *emm*28 isolates carried the A_−27_T_−22_C_−18_ low activity associated promoter but inclusion of international genomic data identified A_−27_T_−22_T_−18_ variant carrying isolates. These two promoter variants were associated with different major lineages within the entire population of 378 international *emm*28 isolates, including one newly sequenced English isolate originally isolated in 1938. The majority of isolates (n=374) clustered either with the reference MGAS6180 strain (USA) (18) or with the reference MEW123 strain (USA) (19) (Figure 7A). Gubbins analysis for core SNP clustering predicted that the two lineages were distinguished by a single 28,200bp region of recombination, between positions 142,426bp (*ntpE*, M28_Spy0126) and 170,625bp (M28_Spy0153) of the MGAS6180 chromosome. This suggests the emergence of one lineage from the other through a single recombination event, followed by expansion of both lineages (Figure 7B). Within the recombination region was the P-*nga-ifs-slo* locus, which differed between the two lineages; although unique in the MGAS6180-like lineage and with low activity associated promoter residues A_−27_T_−22_C_−18_, the MEW123-like lineage had a P-*nga-ifs-slo* identical to that found in *emm*78 isolates (Figure 4), with the three key residues of A_−27_T_−22_T_−18_. This is supported by recent findings identifying two main lineages within *emm*28 and that the A_−27_T_−22_T_−18_ promoter variant conferred greater toxin expression than A_−27_T_−22_C_−18_ (15).

**Figure 7.**
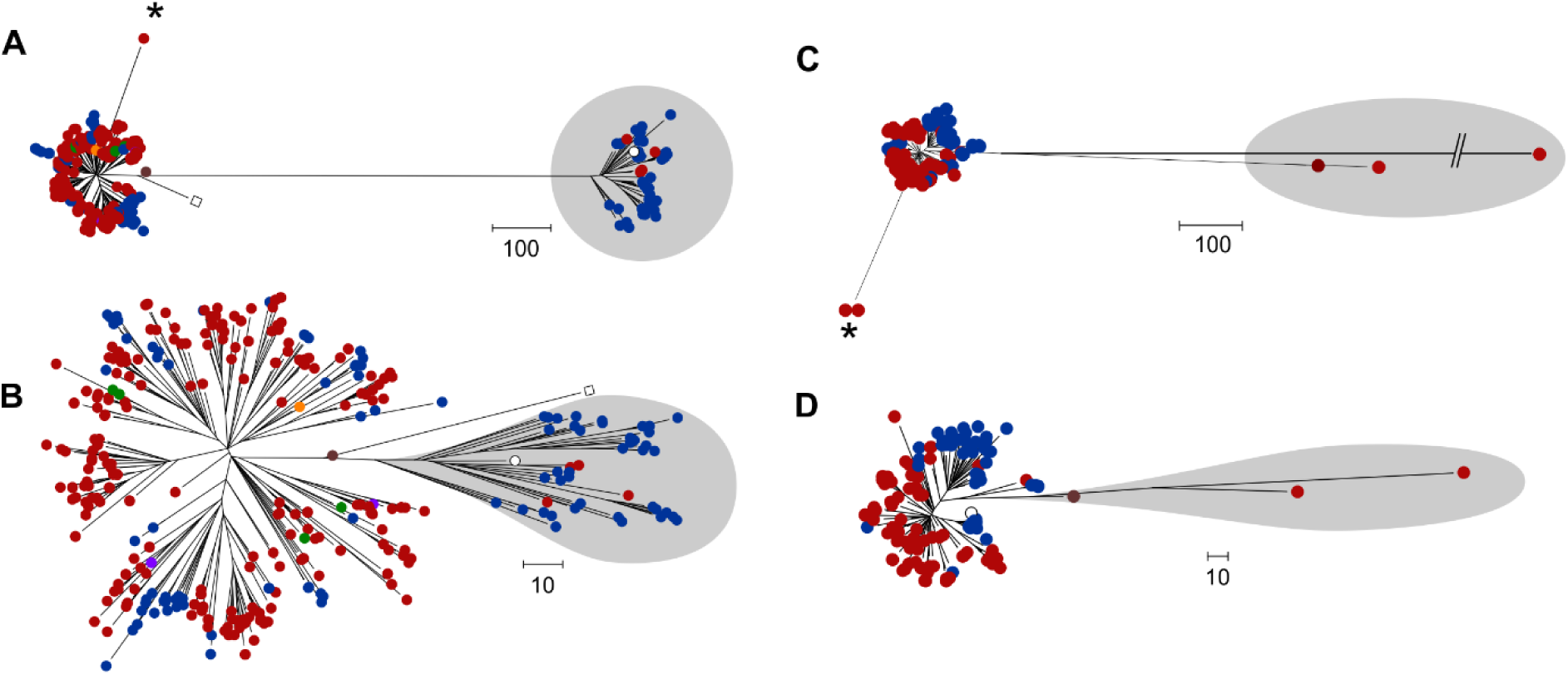
Recombination within the *emm*28 and *emm*87 populations. **(A)** Maximum likelihood phylogeny constructed with core SNPs following mapping of all available *emm*28 genome data to the *emm*28 MGAS6180 reference genome (white square) (18). Modern UK isolates (red circles); BSAC (n=15), CUH (n=13 (11)) and PHE-2014/15 (n=240 (12, 13)), one historical English isolate from 1938 (brown circle). North American isolates (blue circles); ABCs-2015 (n=95 (14)), Canada (2011-2013, n=4 (42)), and completed genome strain HarveyGAS (USA, 2017 (43)). Other isolates; Lebanon (n=1, orange circle (44)), Australia (n=5, green circle (45)), France (STAB10015 (46), M28PF1 (47), purple circles). Two lineages of *emm*28 were identified, one clustering with MGAS6180 (white square) and the other (shaded grey) clustering with MEW123 (2012 USA (19), white circle). (**B)** Regions of recombination were then identified within the *emm*28 genome alignment and removed before reconstructing the phylogenetic tree (**C**) Maximum likelihood phylogeny constructed with core SNPs following mapping of all available *emm*87 genome sequence data to the reference *emm*87 strain NGAS743 (Canada, white circle (48)). UK isolates (red circles); BSAC (2001-2011, n=22), CUH (2008, n=1 (11)), PHE-2014/15 (n=64, (12, 13)). North American isolates (blue circles); ABCs-2015 (n=26, (14)), Canada (n=26, (48)), Texas Children’s Hospital (2012-2016, n=27, (49)). Three isolates (shaded grey) were distinct from the main population. The branch was shortened for one isolate for presentation purposes. (**D**) Regions of recombination were identified within the *emm*87 genome alignment and removed before reconstructing the phylogenetic tree. Isolates indicated by * in both *emm*28 and *emm*87 populations were predicted to have undergone recombination in regions surrounding the *hasABC* locus. Scale bar represents single nucleotide polymorphisms. PHE-2014/15 *emm*28 isolates GASEMM1261, GASEMM2648, GASEMM1396 and GASEMM1353 were removed for presentation purposes as they represented highly divergent lineages.

Although we identified an A_−27_G_−22_T_−18_ high activity variant of P-*nga-ifs-slo* within *emm*28, this was only associated with the highly divergent GASEMM1261 isolate that may represent an *emm* switching event. This isolate, along with three other PHE-2014/15 isolates (GASEMM2648, GASEMM1396 and GASEMM1353) also representing highly divergent lineages, were excluded from the phylogenetic analysis.

All *emm*28 isolates, regardless of lineage and including MGAS6180 (originally isolated in the 1990s), had the same insertion mutation within *hasA* of an A residue after 219 bp. This insertion was predicted to lead to a frameshift and a premature stop codon after 72 amino acids (aa) instead of full length 420 aa, rendering *hasA* a pseudogene. Some isolates also had additional mutations in *hasA*; a deletion of a A residue in a septa-A tract leading to a frameshift and a stop codon after 7 aa (n=1); a deletion of a T residue in a septa-T tract leading to a frameshift and a stop codon after 15 aa (n=2); an insertion of a A residue after 57 bp leading to a frameshift and a stop codon after 46 aa (n=3). The loss of full length HasA would render the isolates acapsular.

In *emm*28 there were just two exceptions where *hasA* found to be intact: the historical *emm*28 isolate from 1938 had an intact *hasABC* capsule operon; and BSAC_bs2099, which appeared to have undergone recombination to acquire a 22,316bp region surrounding the *hasABC* genes, that was 99% identical to the same region in *emm*2 isolate MGAS10270, suggesting *emm*2 might be the donor for this recombination. Both isolates were predicted to express full length HasA and synthesise capsule. Taken together, in comparison with the oldest *emm*28 isolate, the data showed that post 1930s *emm*28 isolates became acapsular through mutation, but the contemporary population is divided into two major lineages, MEW123-like and MGAS6180-like lineages, that may differ in *nga-ifs-slo* expression. Additionally, there was evidence of geographical structure in the population: the MEW123-like lineage comprised mainly of North American isolates (39/44) and only five from England/Wales; isolates from Australia, France and Lebanon were MGAS6180-like, along with the rest of the England/Wales isolates.

Phylogenetic analysis of the BSAC *emm*87 population was expanded and compared with publicly available *emm*87 genome sequence data, totalling 173 isolates from the UK and North America, including one historical NCTC UK isolate from ∼1970-80 (NCTC12065). Gubbins analysis predicted a single 20,506bp region of recombination surrounding the P-*nga-ifs-slo* region, that distinguished the main population from the oldest BSAC isolates from 2001 and the historical 1970-80 NCTC isolate (Figure 7C). Whilst the two 2001 BSAC isolates and the NCTC isolate had a P-*nga-ifs-slo* variant with low activity-associated promoter residues, G_27_T_−22_T_−18_, all other *emm*87 isolates had a P-*nga-ifs-slo* region with high activity associated promoter residues, A_−27_G_−22_T_−18,_ identical to that found in *emm*4 and some *emm*77. This suggested the emergence of a new lineage through a single recombination event followed by expansion within the population, redolent of that previously observed in *emm*89 (Figure 7D).

Similar to *emm*28, all *emm*87 isolates, bar four had an insertion of an A residue after 57 bp that resulted in a frameshift mutation in *hasA,* and the introduction of a premature stop codon after 46aa of HasA. This mutation was also identified within the historical NCTC isolate, but was not found in the two 2001 BSAC isolates, that had an intact *hasABC* locus. This mutation was also absent in two PHE-2014/15 isolates that had undergone an additional recombination event (32,243bp) surrounding the *hasABC* locus, although, as this region shared 100% DNA identity to *emm*28 isolate MGAS6180, HasA is truncated. Overall the data showed that, like *emm*89, contemporary *emm*87 are acapsular with a high activity *nga-ifs-slo* promoter, suggesting that this *emm* lineage may have recently shifted towards this genotype/phenotype.

### Recombination within different multi-locus sequence types of emm75

The *emm*75 genotype is of interest as a common cause of non-invasive infection in the UK; it is also used in models of nasopharyngeal infection (20, 21). Eleven *emm*75 isolates were present in the BSAC collection, all multilocus sequence type (ST) 150. When we incorporated other available genome sequence data for *emm*75 (n=173), including two newly sequenced historical *emm*75 isolates from 1937 and 1938, two major lineages were identified, characterised by two different MLSTs; ST49 or ST150 (Figure 8A). Although the two historic English isolates were ST49, like the majority of modern North American isolates, the modern England/Wales isolates were predominantly ST150.

**Figure 8.**
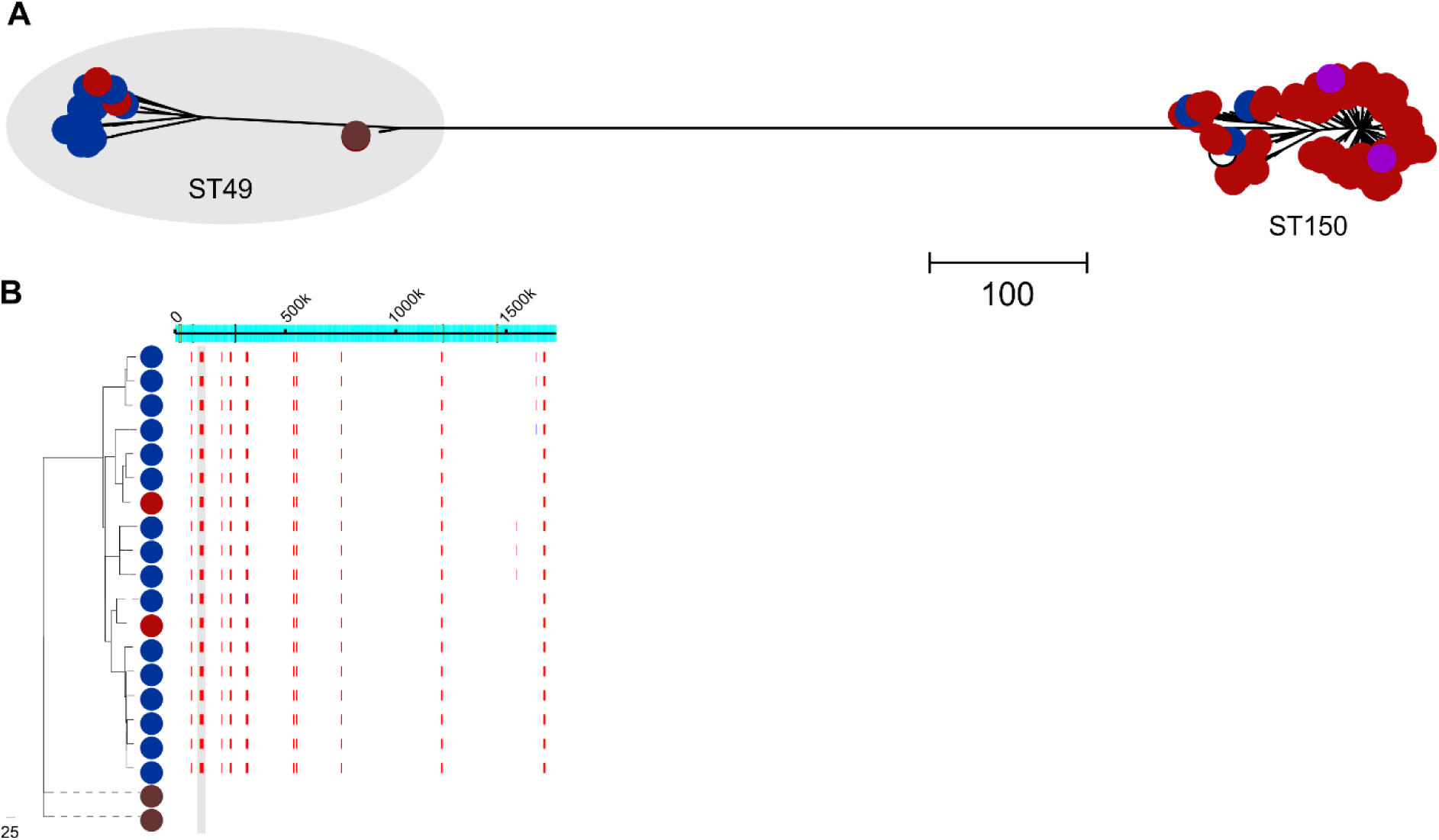
Two lineages within *emm*75. **(A)** Maximum likelihood phylogeny constructed with core SNPs following mapping of all available *emm*75 genome sequence data to the French strain STAB090229 (white circle) (50). Modern UK collections (red circles); BSAC (n=11), CUH (n=6 (11)), PHE-2014/15 (n = 141, (12, 13)) and two English historical isolates (brown circles) from 1937/1938. North American isolates (blue circles); ABCs-2015 (n=20, (14)), NGAS344 and NGAS604 from Canada 2011/2012 (42). French strains (purple circles); STAB120304 (2012) and STAB14018 (2014). Two lineages were identified, generally characterised by the MLST; ST49 (shaded grey) or ST150 (with minor MSLT variants ST788, ST851, ST861 within these lineages). (**B)** Gubbins analysis identified ten regions of predicted recombination (red lines) in all modern ST49 compared to historical 1930s ST49 across the genome (indicated across the top). One region included P-*nga-ifs-slo* (shaded grey). Scale bars represent single nucleotide polymorphisms. One PHE-2014/15 isolates (GASEMM1722) was excluded for presentation purposes as it was highly divergent from the rest of the population.

Although these two ST lineages differed in the P-*nga-ifs-slo* region there was a high level of predicted recombination across the genomes of both STs, perhaps indicative of historic *emm* switching or extensive genetic exchange. ST49 isolates had the subtype 1.1 low activity A_−27_T_−22_C_−18_ promoter, whereas all ST150 isolates had the A_−27_G_−22_T_−18_ subtype 3.1 high activity promoter variant, identical to that of *emm*1/*emm*89 (Figure 4). Modern ST49 isolates did, however, differ from historic 1930s isolates by ten distinct regions of predicted recombination (Figure 8B), including a region spanning the *nga-ifs-slo* locus, although this did not include the promoter region. We did not detect any mutations affecting the capsule region in *emm*75. Taken together, *emm*75 was characterised by two major MLST lineages differing in P-*nga/ifs/slo* promoter activity genotypes but without evidence of recent recombination or loss of capsule.

### Lineages associated with recombination in emm76, emm77 and emm81

The phylogeny of all available genome data for *emm*76, *emm*77 and *emm*81 confirmed the presence of diverse lineages, associated with different MLSTs (Figure 9). In all genotypes, however, there was a dominant MLST lineage representing the majority of isolates; ST50 *emm*76, ST63 *emm*77 and ST624 *emm*81. Within the dominant MLST lineages of *emm*76 and *emm*77, there were sub-lineages that were associated with different P-*nga-ifs-slo* variants as well as loss of functional HasA through mutation.

**Figure 9.**
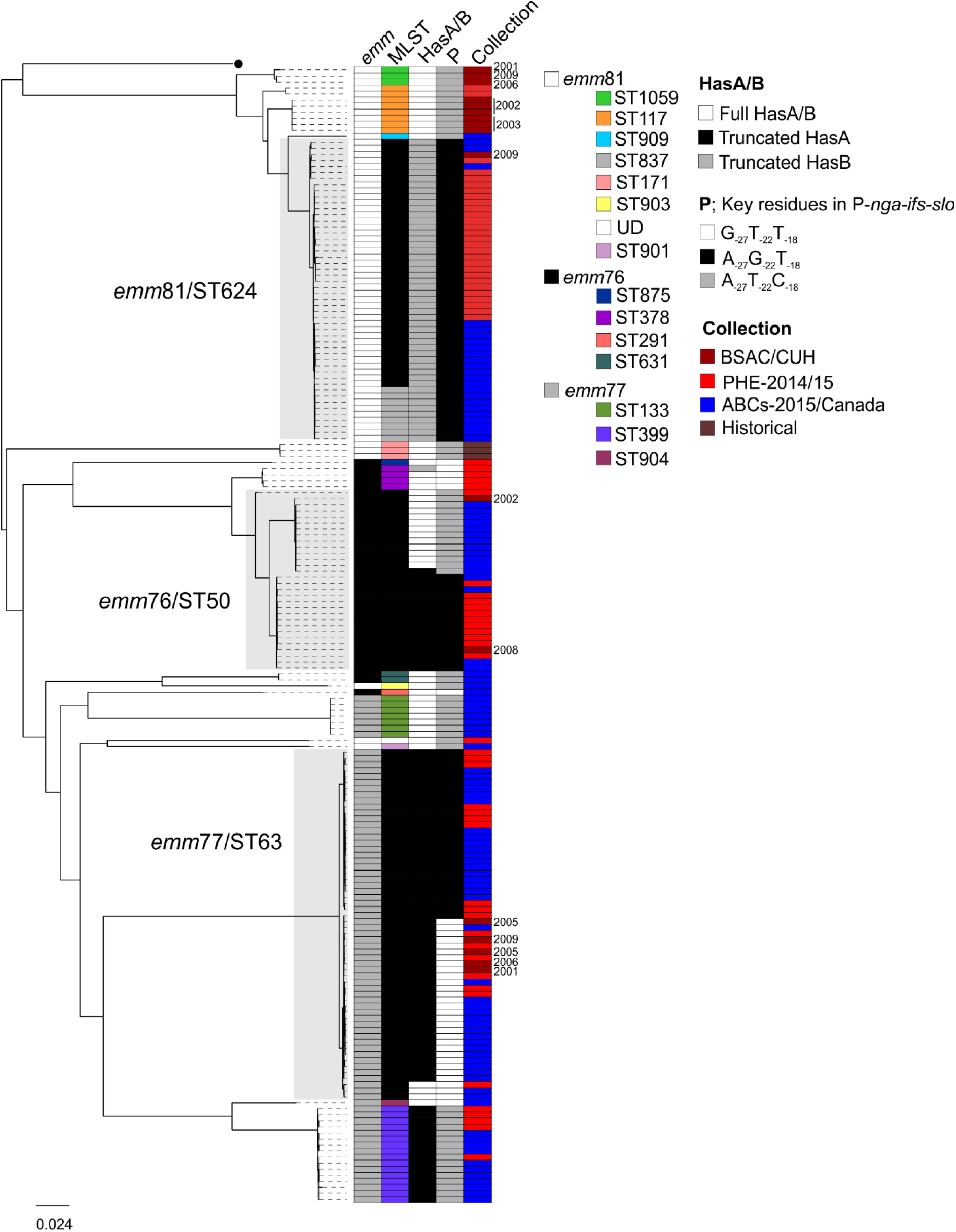
Variants of P-*nga-ifs-slo* and capsule mutations associated with lineages of *emm*76, *emm*77 and *emm*81. Maximum likelihood phylogeny identified multiple MLST lineages within the populations of *emm*76, *emm*77 and *emm*81 (STs provided in the key, UD; undetermined). Major ST lineages are indicated and shaded grey. All *emm*81 isolates were predicted to express full length HasA but the ST624, and minor (single base change in *recP*) ST variant ST837, carry a mutation within *hasB* leading to a truncated HasB. For *emm*76 and *emm*77, mutations were detected in *hasA*. We also identified variants of P-*nga-ifs-slo* associated with one of three combinations of key promoter residues including the high activity associated A_−27_G_−22_T_−18_. Collection indicates either BSAC or CUH (dark red), PHE-2014/15 isolates (red), North America (blue) or English historical (brown). Dates for BSAC isolates and CUH are shown; other isolates were from 2014/2015 or 1930s (historical). *emm*76; n=2 BSAC, n=18 PHE-2014/15 (12, 13), n=18 ABCs-2015 (14). *Emm*77; n=5 BSAC, n=21 PHE-2014/15 (12, 13), n=54 ABCs-2015 (14), n=2 Canada (date unknown) (42). *emm*81; n=9 BSAC, n=1 CUH (11), n=29 PHE-2014/15 (12, 13), n=26 ABCs-2015 (14), n=3 historical 1930s. All sequence data was mapped to the reference strain H293 (black circle). Scale bar represent substitutions per site.

We identified five different MLSTs within *emm*76, but the majority of isolates (30/38) belonged to ST50, including both BSAC isolates. Recombination analysis of the ST50 lineage identified a sub-lineage that differed from other ST50 isolates by 19 regions of recombination (Supplementary Figure 3). One of these regions encompassed P-*nga-ifs-slo*, conferring a P-*nga-ifs-slo* variant closely related to that of modern *emm*1 and *emm*89 with an identical high activity promoter (subtype 3.1). This sub-lineage was dominated by PHE-2014/15 isolates and also contained the more recent of the two BSAC isolates (2008). All isolates in this sub-lineage also had a nonsense mutation within *hasA* of a C to T change at 646bp, resulting in a premature stop codon after 215aa, likely to render the isolates acapsular. Only one ST50 isolate outside this sub-lineage had the same *hasA* C646T change. All other *emm*76 isolates would express full length HasA. This suggests the mutation in *hasA* occurred prior to the recombination events.

Two sub-lineages were also identified within the dominant *emm*77 lineage ST63, and one was associated with the high activity cluster P-*nga-ifs-slo* variant, compared to predicted low activity variants found in the other *emm*77 lineages. Recombination analysis predicted only two regions of recombination distinguishing the two sub-lineages; a region of 17,954bp surrounding P-*nga-ifs-slo*, and a 173bp region within a hypothetical gene (SPYH293_00394) (Supplementary Figure 4). Whilst all BSAC *emm*77 isolates (years 2001-2009) were ST63 with low activity P-*nga-ifs-slo*, PHE isolates from 2014-2015 were almost evenly divided between the two sub-lineages, indicating a potential recent change in England/Wales. All ST63 isolates except three, had a deletion of a T residue within a septa-polyT tract at 458bp in *hasA*, predicted to truncate the HasA protein after 154aa. The three exceptions were predicted to encode full length HasA and were associated with low P-*nga-ifs-slo* promoter activity variants. Although also not associated with high P-*nga-ifs-slo* promoter activity variants, other lineages of *emm*77 also carried mutations within *hasA* that would truncate HasA; ST399 isolates carried an insertion of a T residue at 71 bp of the *hasA* gene resulting in a premature stop codon after 46 aa, and two ST133 isolates carried G894A substitution resulting in a premature stop codon after amino acid residue 297.

The *emm*81 population (n=68) was more diverse with nine different sequence types, but the majority of isolates (41/68) were ST624 or the single locus variant ST837 (9/68; one SNP in *recP* allele) within the same lineage (Figure 9). ST171 was restricted to three historical isolates originally collected in 1938-1939. We did not detect any *hasABC* variations that would disrupt translation in *emm*81 lineages except for the dominant group of ST624/ST837, where we identified an A residue insertion at 128 bp in *hasB* resulting in a frameshift and premature stop codon after 50 aa. All ST624/ST837 carried the high activity cluster P-*nga-ifs-slo* variant identical to that seen in *emm*3, compared to all other lineages associated with other low activity P-*nga-ifs-slo* variants. Recombination analysis identified extensive recombination had occurred within *emm*81 leading to the different levels of diversity, but we identified one region of recombination that distinguished in the ST624/ST837 lineage compared to the closely related ST909 and ST117 populations (Supplementary Figure 5). This region surrounded the P-*nga-ifs-slo* locus, suggesting ST624/ST837 gained the high activity cluster P- *nga-ifs-slo* variant through recombination, like other *emm*-types, potentially from *emm*3 (Figure 4). The prevalence of the high activity and truncated HasB ST624/ST837 lineage may be a recent event in England/Wales, as all BSAC isolates prior to 2009 were outside of this lineage.

### High activity cluster P-nga-ifs-slo variants gained by recombination in emm94 and emm108

Within *emm*94, we identified a P-*nga-ifs-slo* identical to that found in *emm*1 with high activity promoter variant subtype 3.1. Phylogenetic analysis of 51 *emm*94 isolates identified a dominant lineage among England/Wales isolates separate to the single USA isolate and two England/Wales isolates (Supplementary Figure 6), that belonged to ST89. Gubbins analysis predicted 11 regions of recombination in all lineage isolates compared to the three outlying isolates, including one (22,648bp) that encompassed P-*nga-ifs-slo*, transferring a high activity A_−27_G_−22_T_−18_ P-*nga-ifs-slo* variant. All *emm*94 isolates contained an indel within *hasB* compared to the reference (H293); losing 6bp and gaining 13bp between 127-133bp. This variation causes a frameshift and would truncate the HasB protein after 45aa.

We identified a similar high activity cluster P- *nga-ifs-slo* variant within a single *emm*108 genome originating from the USA. Within the 9 isolates from PHE-2014/15 (n=7) and ABCs-2015 (n=2), there were two sequence types, ST1088 and ST14. ST14 was represented by the only two ABCs-2015 isolates and we identified that both had lost the entire *hasB* gene, although *hasA* and *hasC* were still present (Supplementary Figure 7). Additionally, one of the ABCs-2015 isolates had undergone recombination of a single ∼29,683bp region surrounding the P-*nga-ifs-slo,* replacing P-*nga-ifs-slo* for one identical to that found in *emm*3 with high activity promoter variant A_−27_G_−22_T_−18_ subtype 3.1.

## Discussion

The emergence of new, internationally successful lineages of *S. pyogenes* can be driven by recombination-related genome remodelling, as demonstrated by *emm*1 and *emm*89. The transfer of a P-*nga-ifs-slo* region conferring increased expression to the new variant was common to both genotypes. In the case of *emm*89, five other regions of recombination were identified in the emergent variant, one resulting in the loss of the hyaluronic acid capsule. Although potentially all six regions of recombination combined underpinned the success of the emergent *emm*89, we have shown here that recombination of P-*nga-ifs-slo* has occurred in other leading *emm-*types as well as a high frequency of capsule loss through mutation. These data point to an association between genetic change affecting capsule and recombination affecting the P-*nga-ifs-slo* locus, conferring increased production of *nga-ifs-slo*; in some cases (notably *emm*87, *emm*89, and *emm*94) this has further been associated with an apparent fitness advantage and expansion within the population.

A number of genotypes were found to be associated with multiple variants of P-*nga-ifs-slo*. The majority of genotypes had P-*nga-ifs-slo* variants with the low activity promoter associated three key residues variants: G_−27_T_−22_T_−18_ or A_−27_T_−22_C_−18_. Only *emm*1, *emm*3 and *emm*12 were exclusively associated with the high activity A_−27_G_−22_T_−18_ variant. We have shown that the same high activity promoter variant is present in isolates belonging to twelve other *emm* types, notably, *emm*76, *emm*77, *emm*81, *emm*87 and *emm*94, although this is not a consistent feature in these genotypes due to *emm*-switching or recombination. We identified four combination of the three key promoter residues and several subtypes of the 67bp promoter that varied in bases other than those at the −27, −22, and −18 key positions. Although some subtypes were restricted to single genotypes, variation in the −40 base led to the subtype 2.2 of G_−27_T_−22_T_−18_ and subtype 3.2 of A_−27_G_−22_T_−18_. We measured the activity of NADase in representative strains and genotypes of these promoter variants and variation in the −40 base did not impact on the activity conferred by the −27, −22, and −18 bases. Although we predicted the level of *nga* and *slo* expression based on the promoter variant, this may not relate to actual expression given the level of other genetic variation between genotypes. However, our consistent findings of lineages emerging following acquisition of the high activity promoter variant supports the hypothesis that this confers some benefit that may relate to increased toxin expression.

Intriguingly, where we identified an acquisition of the high activity promoter variant through recombination, these genotypes also had a genetic change in the capsule locus, likely rendering the organism unable to make capsule (*hasA* mutation) or only low levels of capsule (*hasB* mutation). To date, only *emm*4, *emm*22, and the emergent *emm*89 lineage are known to lack all three genes required to synthesise capsule. Here, we identified mutations that would truncate HasA and HasB in 35% of all isolates and 65% (35/54) of all genotypes. As the majority of isolates included in this study were invasive or sterile site isolates, the findings further challenge the dogma that the hyaluronan capsule is required for full virulence of *S. pyogenes* and, in addition, lend credence to the possibility that the increased expression of NADase and SLO may in some way compensate for the lack of capsule (22). While capsule has been shown to underpin resistance to opsonophagocytic killing in the most constitutively hyper-encapsulated genotypes such as *emm*18 (23, 24), there is less evidence that it contributes measurably to opsonophagocytosis killing resistance in other genotypes (3). Whether loss of capsule synthesis is of benefit to *S. pyogenes* is uncertain; the capsule may shield several key adhesins used for interaction with host epithelium and fomites, but may also act as a barrier to transformation with DNA. An accumulation of *hasABC* inactivating mutations have been identified during long term carriage (25) and, although for some genotypes capsule loss impacted on survival in whole human blood, a high number of acapsular *hasA* mutants have also recently been found to be causing a high level of disease in children, including *emm*1, *emm*3 and *emm*12 (26).

The process of recombination in *S. pyogenes* is not well understood and natural competence has only been demonstrated once and under conditions of biofilm or nasopharyngeal infection (27). We do not know if the six regions of recombination that lead to the emergence of the new ST101 *emm*89 variant occurred simultaneously, although no intermediate isolates have been identified. The loss of the hyaluronic acid capsule in the new emergent *emm*89, along with our consistent findings of inactivating mutations associated with P-*nga-ifs-slo* transfer indicate either 1) the process of recombination requires the inactivation of capsule, 2) capsule negative *S. pyogenes* requires high expression of *nga-ifs-slo* for survival, 3) or that capsule negative phenotype combined with high expression of *nga-ifs-slo* provides a greater selective advantage to *S. pyogenes*.

The phylogeny of *emm*28, *emm*87, *emm*76, *emm*77, *emm*94, and *emm*108 indicated that mutations in *hasA* or *hasB* occurred prior to recombination of P-*nga-ifs-slo*, supporting the first hypothesis that prior capsule inactivation is required for recombination. There is no evidence, however, to suggest this was required for recombination in the *emm*1 population. It could be hypothesised that capsule acts as a barrier to genetic exchange, but there has also been a positive genetic association of capsule to recombination rates (28). A positive association may, however, be related only to species expressing antigenic capsule whereby recombination is required to introduce variation for immune escape.

The *hasC* gene is not essential for capsule synthesis (29) because a paralog of *hasC* exists within the *S. pyogenes* genome. A paralog for *hasB* (*hasB.2*) also exists elsewhere in the *S. pyogenes* chromosome and can act in the absence of *hasB* to produce low levels capsule (30) but *hasA* is absolutely essential for capsule synthesis (29). The mutations in *hasA* in *emm*28 and *emm*87 have been previously noted and confirmed to render the isolates acapsular (26, 31). Not all acapsular isolates were found to carry the high activity promoter of *nga-ifs-slo*, despite being invasive, perhaps refuting the hypothesis that high activity *nga*-*ifs-slo* promoter is essential for the survival of acapsular *S. pyogenes*.

Interestingly, we identified that the capsule locus is also a target for recombination as, like *emm*89, isolates within *emm*28 and *emm*87 had undergone recombination of this locus and surrounding regions, varying in length (Supplementary Figure 8) and restoring capsule synthesis in *emm*28. Isolated examples of loss of *hasA* or *hasB* genes were identified in some genotypes, such as *emm*108, possibly due to internal recombination and deletion.

Only two *emm*4 and one *emm*22 isolates were found to have P-*nga-ifs-slo* variants that were not an A_−27_T_−22_G_−18_ high activity promoter variants, and interestingly these isolates carried the *hasABC* genes, typically absent in *emm*4 and *emm*22. The high genetic distance of these isolates to the other *emm*4 and *emm*22 genomes indicated potentially *emm* switching of the *emm*4 or *emm*22 genes onto different genetic backgrounds. The single *emm*28 with a high activity P-*nga-ifs-slo* variant also may be an example of this, and was one of four *emm*28 isolates that did not cluster with the two main *emm*28 lineages. Although we excluded them from our analysis as we focussed on recombination within the two main lineages, this potential for highly diverse variants within genotypes and the potential for *emm*-switching warrants further investigation, particularly as the most promising current vaccine is multi-valent towards common M types (32).

All other genotypes carrying the high activity P-*nga-ifs-slo* variant were found to have undergone recombination of this region; *emm*28, *emm*75, *emm*76, *emm*77, *emm*81, *emm*87, *emm*94 and *emm*108, as well as the previously described *emm*1 and *emm*89.

Within *emm*87, we identified three isolates outside of the main population lineage that represented the oldest isolates in the collection; two from 2001 (different geographical locations within England) and one from ∼1970-80. A single region of recombination, surrounding the P-*nga-ifs-slo* locus distinguished the main population lineage from the three older isolates, consistent with a recombination event but, due to a lack of earlier isolates of *emm*87, we could not confirm a recombination related shift in the population, as reported previously for *emm*89 and *emm*1.

The existence of two lineages within the contemporary *emm*28 suggests that one has not yet displaced the other, although the MEW123-like lineage was predominantly USA isolates, consistent with recent findings (15). The P-*nga-ifs-slo* region with the high activity associated A_−27_T_−22_T_−18_ and acquired through recombination by the MEW123-like lineage was identical to that found in *emm*78, indicating *emm*78 as the potential genetic donor. We found *emm*78 to have high levels of NADase activity, as predicted, and interestingly, like *emm*28, all eight *emm*78 isolates were acapsular due to a deletion within the *hasABC* promoter region extending into *hasA*. This again may support the hypothesis that capsule negative *S. pyogenes* requires high expression of *nga-ifs-slo* for survival.

A strength of this study was the systematic longitudinal sampling over a 10 year period; as expected, this again identified the shift in the *emm*89 population. Other *emm*-types exhibited lineages with different P-*nga-ifs-slo* variants, and those with the more active promoter variant did appear to become dominant over time, similar to *emm*1 and the emergent *emm*89 lineages. For example, the high activity P-*nga-ifs-slo* ST63 lineage of *emm*77 was not detected in England/Wales isolates prior to 2014-15. Similarly, the high activity P-*nga-ifs-slo* variant *emm*81 ST646/ST837 lineage was represented by only a single isolate (of six) collected 2001-2009 but became dominant by 2014/15 in England/Wales and the USA. *emm*75 was the 6^th^ most common genotype in England/Wales 2014-15 and dominated by high activity P-*nga-ifs-slo* variant ST150 lineage, yet less common in the USA where ST49 with low activity P-*nga-ifs-slo* is dominant. A high prevalence of *emm*94 was also found in England/Wales 2014-15 but was rare in the USA (only 1 isolate). Our analysis of this genotype indicated there has been a recombination related change in the population as we detected 11 regions of predicted recombination including P-*nga-ifs-slo* potentially conferring high toxin expression. The other ten regions of recombination may also provide advantages to this lineage along with a potential low level of capsule through *hasB* mutation.

The development and boosting of circulating antibodies to SLO is often used as a diagnostic biomarker of recent *S. pyogenes* infection and is known to be more specific to throat rather than skin infections. The genomic analysis provides explanation for this historic and well-recognized association between anti-SLO titres and disease patterns, due to known tissue tropism of *S. pyogenes emm* types. Whether the alteration of SLO activity in different *S. pyogenes* strains might render such a test more or less specific will be of interest, although may explain observed differences in ASO titre between genotypes (33). There is also the possibility that other beta haemolytic streptococci might acquire similarly active SLO production, reducing the specificity of ASO titre to *S. pyogenes*.

Our genomic analysis has uncovered convergent evolutionary pathways towards capsule loss and recombination related re-modelling of the P-*nga-ifs-slo* locus in leading contemporary genotypes. This suggests that a combination of capsule loss and gain of high *nga-ifs-slo* expression provides a greater selective advantage than either of these phenotypes alone. Acquisition of the high activity promoter led to pandemic *emm*1 and *emm*89 clones that are dominant and highly successful. Active surveillance of the lineages comprising *emm*76, *emm*77, *emm*81, *emm*87, *emm*94 and *emm*108 is required to determine if capsule loss/reduction and recombination of P-*nga-ifs-slo* towards high expression will trigger expansion towards additional pandemic clones in the next few years.

## Materials & Methods

### Isolates

344 isolates of *S. pyogenes* associated with blood stream infections and submitted to the British Society for Antimicrobial Chemotherapy (BSAC, www.bsacsurv.org) from 11 different sites across the UK between 2001-2011 were subjected to whole genome sequencing (Supplementary Table 1). All BSAC isolates were tested for antibiotic susceptibility using the BSAC agar dilution method to determine MICs (34).

A further six isolates were sequenced from a historical collection of *S. pyogenes* originally collected in the 1930s from puerperal sepsis patients at Queen Charlottes Hospital, London, UK; one *emm*28 from 1938 (ERR485803), two *emm*75 from 1937 (ERR485807) and 1939 (ERR485820), three *emm*81 from 1938 (ERR485805) and 1939 (ERR485801, ERR485802).

### Genome sequencing

Streptococcal DNA was extracted using the QIAxtractor instrument according to the manufacturer’s instructions (QIAgen, Hilden, Germany), or manually using a phenol-chloroform method (35). DNA library preparation was conducted according to the Illumina protocol and sequencing was performed on an Illumina HiSeq 2000 with 100 - cycle paired-end runs. Sequence data have been submitted to the European Nucleotide Archive (ENA) (www.ebi.ac.uk/ena) (accession numbers in Supplementary Table 2).

Genomes were *de novo* assembled using Velvet with the pipeline and improvements found at https://github.com/sanger-pathogens/vr-codebase and https://github.com/sanger-pathogens/assembly_improvement (36). Annotation was performed using Prokka. *emm* genotypes were determined from the assemblies and multilocus sequence types (MLSTs) were identified using the MLST database (pubmlst.org/spyogenes) and an in-house script (https://github.com/sanger-pathogens/mlst_check). New MLST were submitted to the database (https://pubmlst.org/). Antimicrobial resistance genes were identified by srst2 (37) using the ARG-ANNOT database (ARGannot_r2.fasta) (38).

### Genome sequence analysis

Sequence reads were mapped using SMALT (https://www.sanger.ac.uk/science/tools/smalt) to the completed *emm*89 reference genome H293 (3) as this genome contains no known prophage regions. Other reference genomes were also used where indicated with predicted prophage regions (Supplementary Table 3) excluded to obtain ‘core’ SNPs. Maximum-likelihood phylogenetic trees were generated from aligned core SNPs using RAxML (39) with the GTR substitution model (39) and 100 bootstraps. Regions of recombination were predicted using Gubbins analysis using the default parameters (40).

Other genome sequence data were obtained from the short read archive. We combined data collected across England and Wales through Public Health England during 2014 and 2015 (PHE-2014/15) supplied by Kapatai *et al*. (13) and Chalker *et al*. (12) from invasive and non-invasive *S. pyogenes* isolates. We also used data supplied by Chochua et al. (14) collected by Active Bacterial Core Surveillance USA in 2015 (ABCs-2015) from invasive *S. pyogenes* isolates. ABCs-2015 sequence data was pre-processed by Trimmomatic (41) to remove adapters and low quality sequences. PHE-2014/15 had already been pre-processed (12, 13). Genome data from these collections were assembled *de novo* using Velvet (assembly statistics provided in Supplementary Table 2) and any isolates with atypical total assembled length or contig numbers were excluded. We also used data from Turner *et al*. (2017) of invasive and non-invasive isolates from the Cambridgeshire region, UK and collected through Cambridge University Hospital (CUH) (11). We relied on the *emm*-type determined during the original studies and excluded any data where the *emm*-type was uncertain or negative. The genes *hasA, hasB, hasC* and the P-*nga-ifs-slo* were extracted from the assembled genome using *in silico* PCR (https://github.com/simonrharris/in_silico_pcr). Capsule locus and P-*nga-ifs-slo* variants were also confirmed through mapping.

### NADase activity

Activity of NADase was measured in culture supernatant as previously described (3). Activity was determined as the highest dilution capable of hydrolysing NAD+.

## Acknowledgments

This publication presents independent research supported by the Health Innovation Challenge Fund (WT098600, HICF-T5-342), a parallel funding partnership between the Department of Health and Wellcome Trust. The work was also funded by the UK Clinical Research Collaboration (UKCRC, National centre for Infection Prevention & Management) and the National Institute for Health Research Biomedical Research Centre awarded to Imperial College London. The views expressed in this publication are those of the author(s) and not necessarily those of the Department of Health, NIHR, or Wellcome Trust. CET was an Imperial College Junior Research Fellow and is a Royal Society & Wellcome Trust Sir Henry Dale Fellow (208765/Z/17/Z).

## Supplementary Figures

**Supplementary Figure 1.**
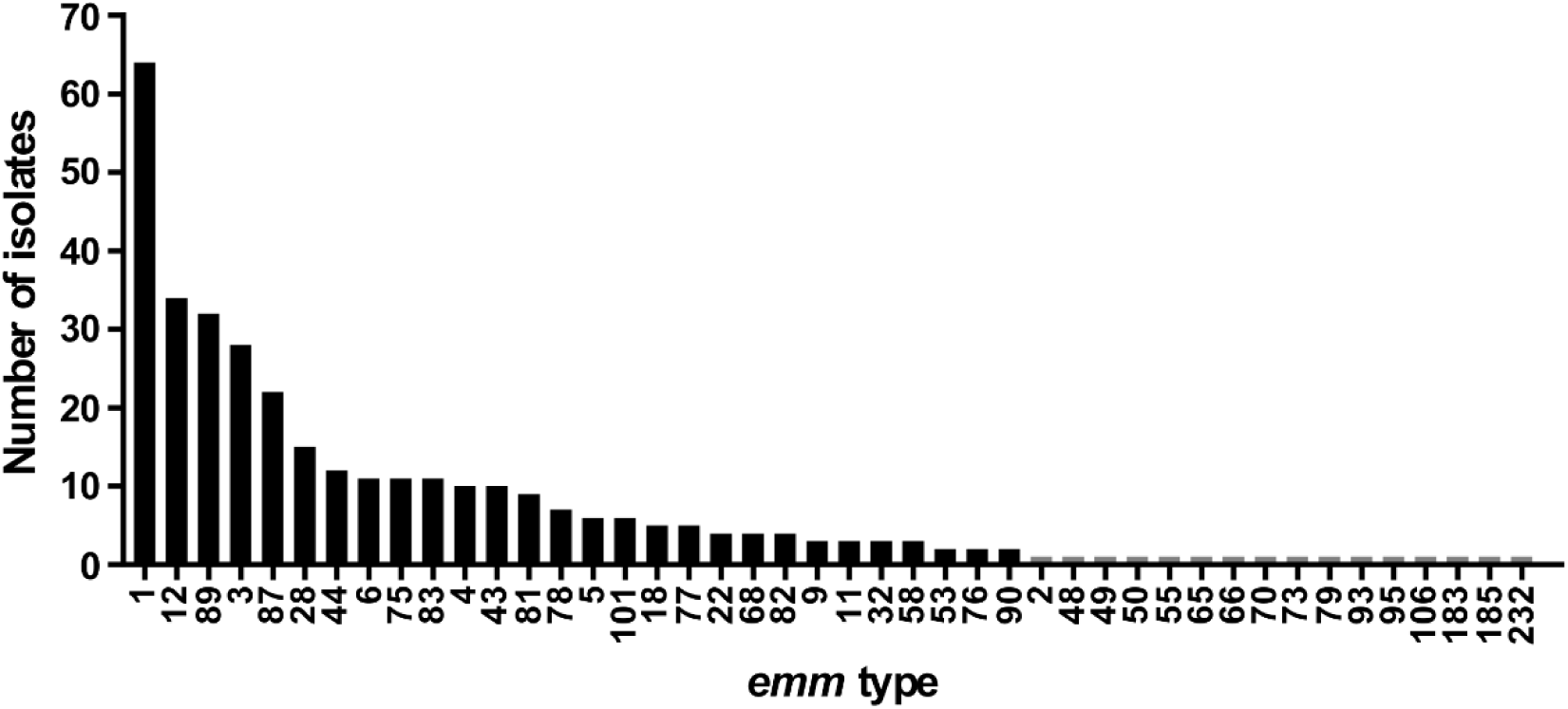
Number of isolates per *emm*-type in the BSAC collection. Forty-four different genotypes were identified within the collection but 16 were represented by single isolates (grey bars). Total number of isolates was 344.

**Supplementary Figure 2.**
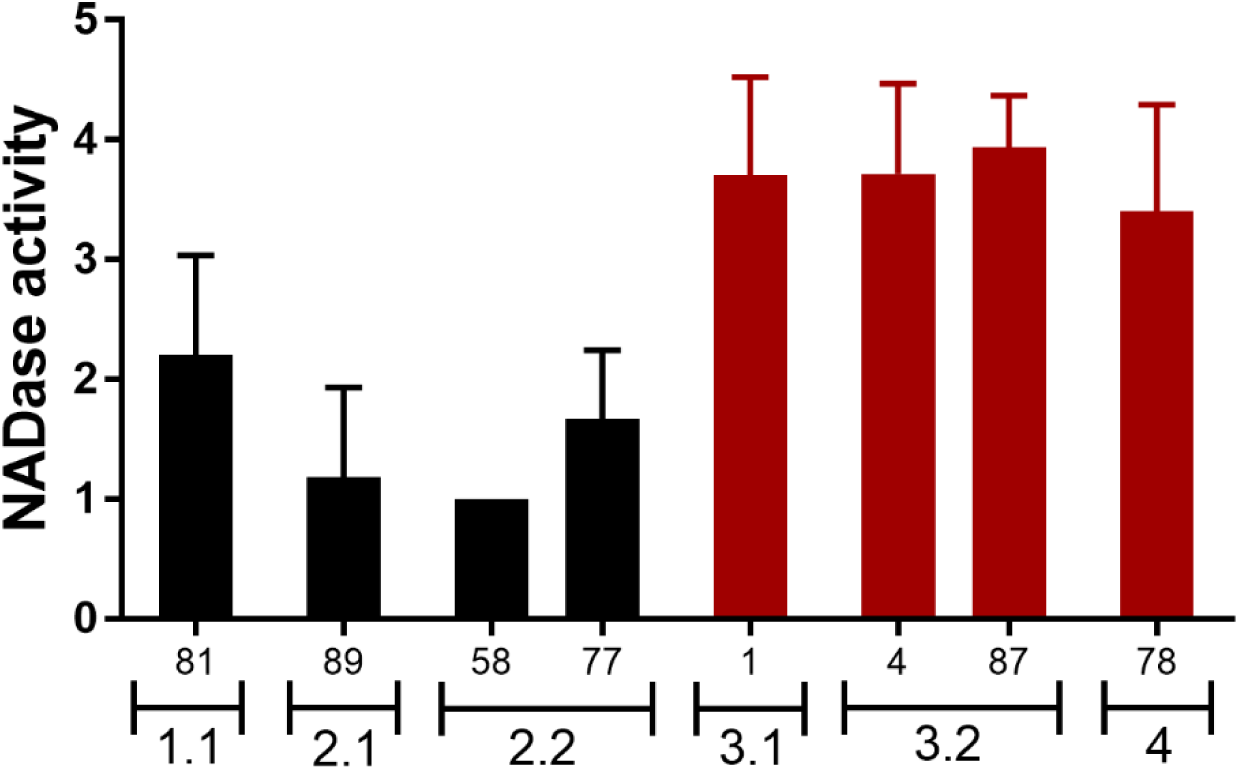
NADase activity of different promoter subtypes. The activity of NADase was measured in culture supernatant of BSAC isolates representing different promoter subtypes with predicted low (black) or high (red) activity. A_−27_T_−22_C_−18_ subtype 1.1 promoter has low activity in *emm*81 isolates, consistent with previous findings of this promoter in historical *emm*1. G_−27_T_−22_T_−18_ subtype 2.1 had low activity in older *emm*89, also consistent with previous findings, and subtype 2.2 in *emm*58 and *emm*77 also had low activity, as predicted despite the additional base change at −40bp. High activity of A_−27_G_−22_T_−18_ subtype 3.1 was confirmed in *emm*1 and subtype 3.2 in *emm*4 and *emm*87 also had high activity, also supporting a null effect of the base change at −40bp. A_−27_T_−22_T_−18_ subtype 4 promoter in *emm*78 had high activity. Isolates with mutations in regulators *covR*/S or *rocA* were excluded as they influence the expression of *nga*. Data represent mean +SD of e*mm*1; n=10, *emm*89; n=11, *emm*58; n=3, *emm*77; n=3, *emm*4; n=7, *emm*87; n=17, *emm*78; n=5, *emm*81; n=5.

**Supplementary Figure 3.**
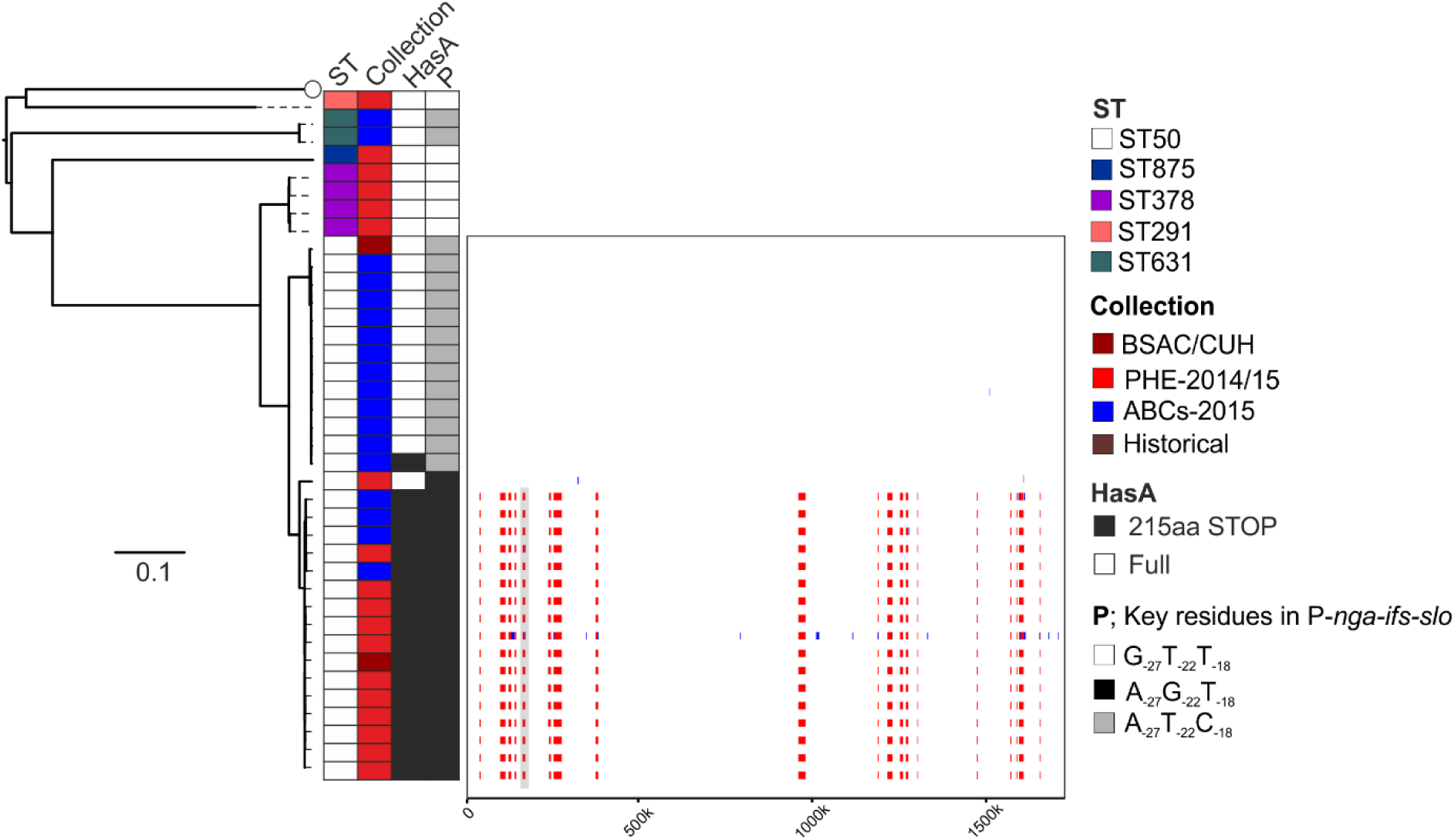
Recombination within ST50 *emm*76. All sequence data for *emm*76 (n=38) were mapped to the reference strain H293 (white circle). Majority of isolates were ST50 and within this ST were two sub-lineages. Recombination analysis (boxed region) of ST50 isolates identified 19 regions of recombination across the genome in all isolates (red vertical lines) belonging to the lower sub-lineage compared to the top sub-lineage. One of these regions (highlighted grey) surrounded the P-*nfa-ifs-slo* locus conferring the high activity associated promoter with residues A_−27_G_−22_T_−18_ to the lower sub-lineage compared to low activity A_−27_T_−22_C_−18_ in the top sub-lineage. Scale bar represents substitutions per site. Scale on boxed region represents position in the H293 genome.

**Supplementary Figure 4.**
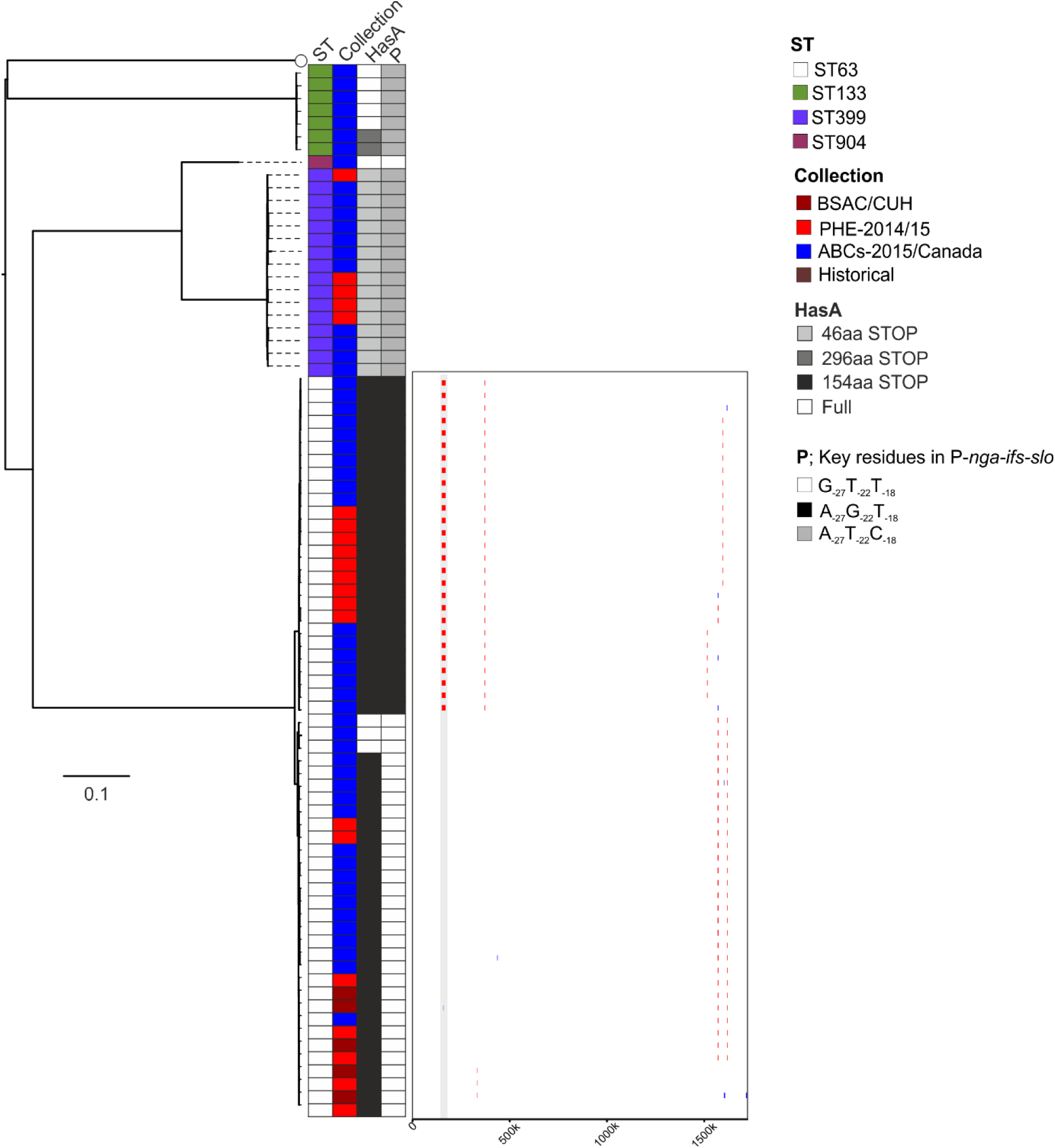
Recombination within ST63 *emm*77. All sequence data for *emm*77 (n=82) were mapped to the reference strain H293 (white circle). Majority of isolates were ST63 and within this ST were two sub-lineages. Recombination analysis (boxed region) of ST63 isolates identified 2 regions of recombination across the genome in all isolates (red vertical lines) belonging to the top sub-lineage compared to the lower sub-lineage. One of these regions (highlighted grey) surrounded the P-*nfa-ifs-slo* locus conferring the high activity associated promoter with residues A_−27_G_−22_T_−18_ to the lower sub-lineage compared to low activity G_−27_T_−22_T_−18_ in the top sub-lineage. Scale bar represents substitutions per site. Scale on boxed region represents position in the H293 genome.

**Supplementary Figure 5.**
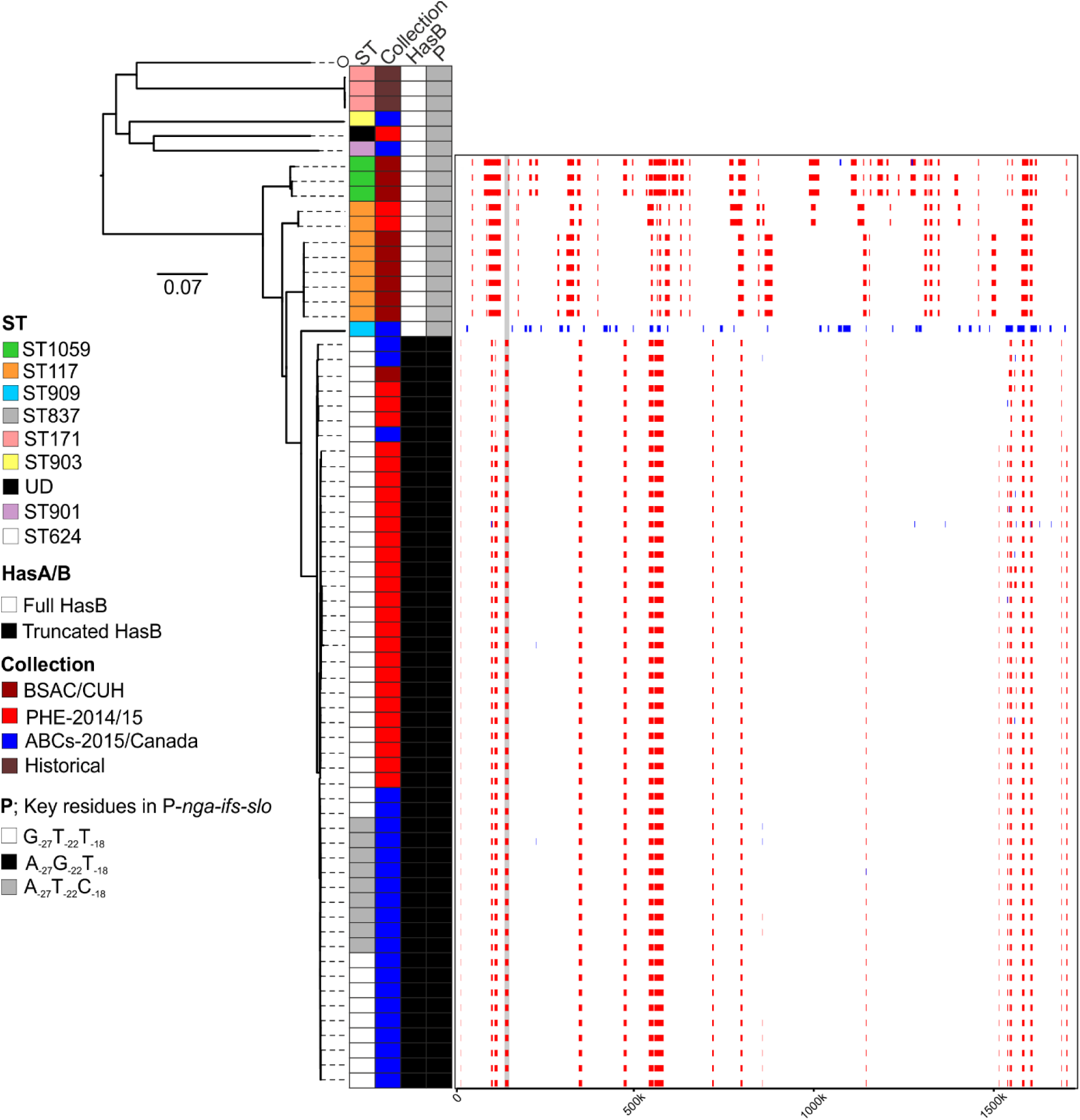
Recombination within *emm*81. All sequence data for *emm*81 (n=68) were mapped to the reference strain H293 (white circle). Majority of isolates were ST624. Recombination analysis (boxed region) of ST624 isolates and closely related ST1059, ST117, ST909 and ST837 identified patterns of recombination across the genome in all isolates (red vertical lines, or blue vertical lines if unique to a single isolate). One of these regions (highlighted grey) surrounded the P-*nfa-ifs-slo* locus conferring the high activity associated promoter with residues A_−27_G_−22_T_−18_ to the ST624/ST837 population compared to low activity G_−27_T_−22_T_−18_ in all other isolates. Scale bar represents substitutions per site. Scale on boxed region represents position in the H293 genome.

**Supplementary Figure 6.**
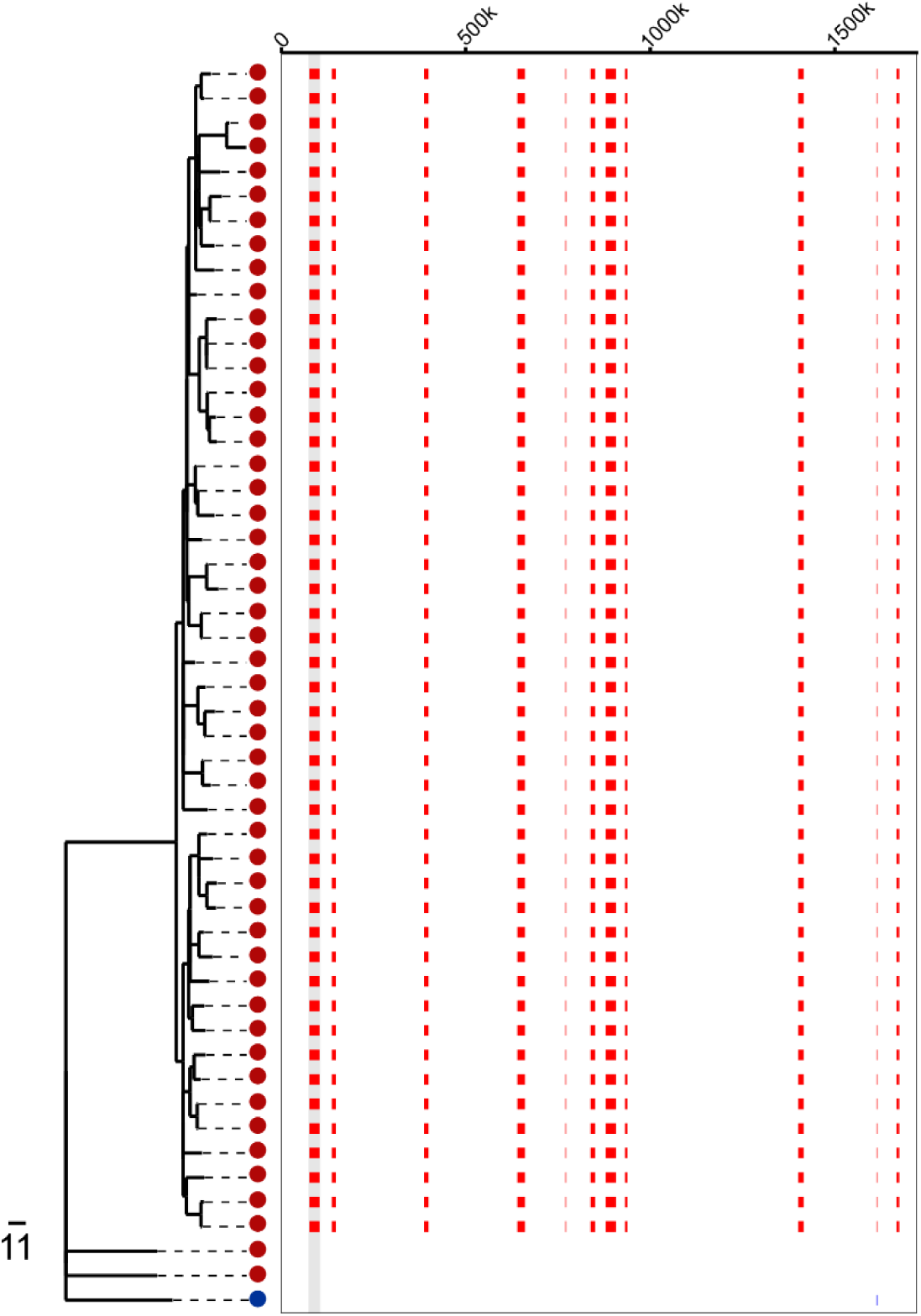
Recombination in *emm*94. In the 2014/2015 UK *emm*94 population, the majority (n=51) form a lineage separate from two 2014/2015 UK isolates and the single 2015 USA isolate. SNP clustering analysis predicted 11 regions of recombination (red lines) in all the lineage associated isolates compared to the three other isolates. One of these regions (highlighted in grey) encompassed the P-*nga-ifs-slo* region. Scale bar represents SNPs.

**Supplementary Figure 7.**
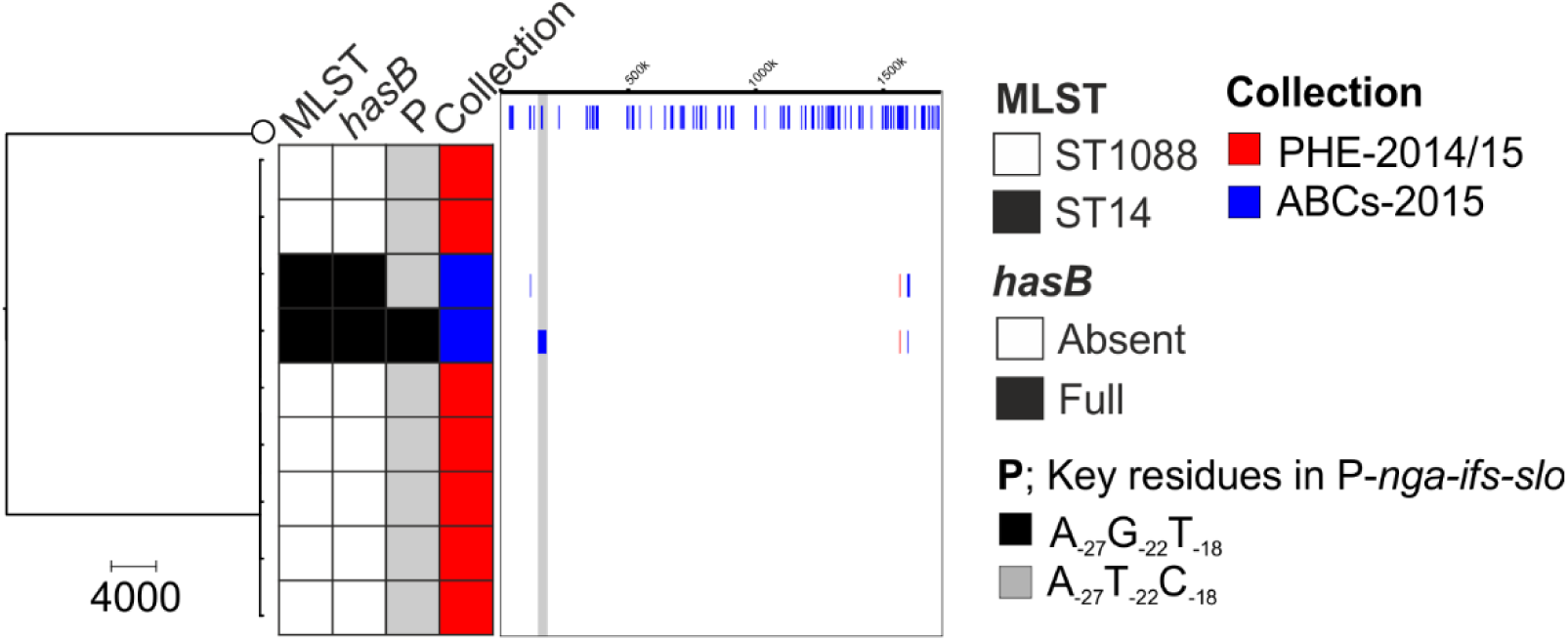
Recombination in *emm*108 around P-*nga-ifs-slo*. Isolates of *emm*108 from the ABCs-2015 (blue) collection were of a different MLST (ST14) compared to PHE-2014/15 (ST1088). The *hasB* gene was absent in the genomes of both ABCs-2015 isolates and one had undergone recombination surrounding the P-*nga-ifs-slo* locus (shaded grey), as predicted by SNP cluster analysis (shown on the right). Blue lines; predicted recombination unique to a single genome. Sequence data were mapped to the reference strain H293, also used as an outgroup for SNP cluster analysis. Scale bar represents SNPs.

**Supplementary Figure 8.**
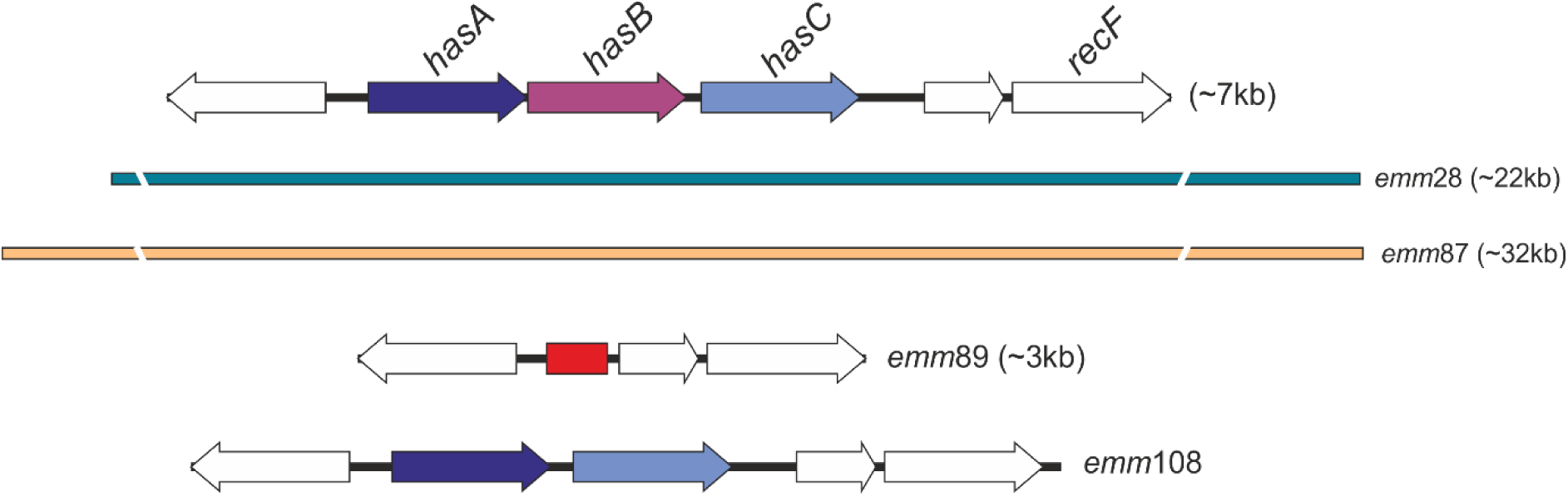
Regions of recombination spanning the capsule locus. Recombination across the *hasA, hasB* and *hasC* genes was identified in two genotypes in addition to the previously described *emm*89. Length of recombination, predicted by SNP cluster analysis, ranges from ∼3kb to 32kb. Recombination within *emm*89 resulted in the loss of all three genes and the gain of a 150bp region (red). In *emm*108, the *hasB* gene was lost but this may be through recombination within the chromosome rather than recombination. All regions are shown relative to the reference genome H293 and genes within this region are depicted as arrows. Recombination in *emm*28 and *emm*87 extended beyond the region depicted and shown as broken lines.

## Supplementary Tables

**Supplementary Table 1 – Details of BSAC isolates and antimicrobial sensitivity testing**

Excel File- Supplementary_Table_1.xlsx

**Supplementary Table 2 – Details of all isolates with assembly statistics, capsule gene mutations and *nga/ifs/slo* promoter variants.**

Excel File – Supplementary_Table_2.xlsx

**Supplementary Table 3.** Reference genomes used for mapping to in this study and excluded prophage regions.

**excluded prophage regions**
| Reference strain | *emm* type | Prophage locations |
| --- | --- | --- |
| H293 | 89 | None |
| MGAS6180 | 28 | 986212-1032479<br>1226909-1269223<br>1807177-1821524<br>1845845-1857621<br>1081040-1092149<br>1286346-1322672 |
| STAB090229 | 75 | 723790-761364<br>1138154-1180947<br>1473588-1512859 |
| NGAS743 | 87 | 547772-585345<br>709624-756401<br>1199244-1242181<br>1253797-1293056 |

